# Extensive sequence divergence of non-coding regions between *Aspergillus fumigatus*, a major fungal pathogen of humans, and its relatives

**DOI:** 10.1101/2021.10.26.465918

**Authors:** Alec Brown, Matthew E. Mead, Jacob L. Steenwyk, Gustavo H. Goldman, Antonis Rokas

**Author notes:** Address correspondence to: Antonis Rokas.

## Abstract

Invasive aspergillosis is a deadly fungal disease; more than 400,000 patients are infected worldwide each year and the mortality rate can be as high as 50-95%. Of the ∼450 species in the genus *Aspergillus* only a few are known to be clinically relevant, with the major pathogen *Aspergillus fumigatus* being responsible for ∼50% of all invasive mold infections. Genomic comparisons of *A. fumigatus* to other *Aspergillus* species have historically focused on protein-coding regions. However, most *A. fumigatus* genes, including those that modulate its virulence, are also present in non-pathogenic close relatives of *A. fumigatus*. Our hypothesis is that differential gene regulation – mediated through the non-coding regions upstream of genes’ transcription start sites – contributes to *A. fumigatus* pathogenicity. To begin testing this, we compared non-coding regions up to 500 base pairs upstream of the first codon of single-copy orthologous genes from the two *A. fumigatus* reference strains Af293 and A1163 and eight closely related *Aspergillus* section *Fumigati* species. We found that non-coding regions showed extensive sequence variation and lack of homology across species. By examining the evolutionary rates of both protein-coding and non-coding regions in a subset of orthologous genes with highly conserved non-coding regions across the phylogeny, we identified 418 genes, including 25 genes known to modulate *A. fumigatus* virulence, whose non-coding regions exhibit a different rate of evolution in *A. fumigatus*. Examination of sequence alignments of these non-coding regions revealed numerous instances of insertions, deletions, and other types of mutations of at least a few nucleotides in *A. fumigatus* compared to its close relatives. These results show that closely related *Aspergillus* species that vary greatly in their pathogenicity exhibit extensive non-coding sequence variation and identify numerous changes in non-coding regions of *A. fumigatus* genes known to contribute to virulence.

## Introduction

Invasive aspergillosis (IA), a human disease caused by members of the fungal genus *Aspergillus*, is responsible for >400,000 cases worldwide per year with a mortality rate between 50-95% (Bongomin et al., 2017). More than 90% of IA cases are caused by *Aspergillus fumigatus*, with about a dozen other species such as *Aspergillus lentulus*, *Aspergillus thermomutatus*, and *Aspergillus udagawae* accounting for the rest (Steinbach et al., 2012; Rokas et al., 2020). Studies in both environmental (Flores et al., 2013) and hospital settings (Wirmann et al., 2018) show that asexual spores (conidia) of *A. fumigatus* and many other *Aspergillus* species are present, yet *A. fumigatus* causes IA more frequently than its close relatives.

IA begins with inhalation of *Aspergillus* asexual spores and subsequent interaction between the asexual spores and the epithelium of the lung (Chotirmall et al., 2013). Several defense mechanisms including physical removal of asexual spores (Croft et al., 2016), secretion of antimicrobial peptides (Wiesner et al., 2017) and recruitment of specialized immune cells are employed by the human host to prevent spore growth (Bertuzzi et al., 2018). To cause infection, *A. fumigatus* must overcome these challenges and adapt to the host environment to survive. The dynamics and intricacies of the interaction between *A. fumigatus* and host responses have yet to be fully elucidated. Decades of work have identified at least 206 genetic determinants of *A. fumigatus* virulence, that is genes whose deletion is known to modulate the virulence of *A. fumigatus* (for a detailed list, see: Steenwyk et al., 2021). These genetic determinants of virulence are involved in a wide range of activities including gene regulation, RNA processing, protein modification, production of secondary metabolites, amino acid biosynthesis, cell cycle regulation, morphological regulation, and others (Steenwyk et al., 2021).

The phylogeny of the genus *Aspergillus* reveals that pathogenic species are often more closely related to nonpathogenic species than to other pathogenic ones (Hourbraken et al., 2014; de Vires et al., 2017; Steenwyk et sl., 2019; Rokas et al., 2020; Mead & Steenwyk et al., 2021). For example, *A. fischeri* is a close relative of *A. fumigatus* (the two share >90% average nucleotide sequence similarity and >95% amino acid sequence similarity between orthologs), yet *A. fischeri* is less virulent and is not considered clinically relevant (Mead et al., 2019; Steenwyk et al., 2020). Given the large disparity of IA cases caused by *A. fumigatus* and closely related species, early studies looked to species-specific genes in *A. fumigatus* as a potential contributor (Fedorova et al., 2008). However, a recent examination found that 206 known genetic determinants of virulence in *A. fumigatus* are shared between *A. fumigatus* and at least one other closely related species (Mead & Steenwyk et al., 2021).

Variation in non-coding regions can also contribute to phenotypic diversity (Carroll, 2008; Li & Johnson, 2012) and disease (Caron et al., 2019; Ropero et al., 2017; Jang et al., 2018). In fungi, non-coding regions are bound by transcription factors (TFs), impact transcriptional activity (Kim et al., 2019), and play roles in vital biological processes such as zinc homeostasis (Eide, 2020) and thermotolerance (Yamamoto et al., 2020). Differences in gene expression have become an important focus in understanding *A. fumigatus* virulence (de Castro et al., 2014, Chung et al., 2014; Furukawa et al., 2020; Ries et al., 2020; Colabardini et al., 2021; Takahashi et al., 2021). However, the role non-coding regions play in differential gene regulation between *A. fumigatus* and close relatives remains a mystery.

Here, we perform genome-wide comparisons of intergenic, non-coding regions of the reference strains *A. fumigatus* Af293 and A1163 against those of eight closely related species. We identified 5,215 single-copy orthologous genes across the 10 taxa of interest. Of the 5,215 genes, 4,483 of their intergenic, non-coding regions either lacked homology across the ten taxa or showed extensive sequence variation, such that multiple sequence alignment was not reliable. Only 732 were of the same length, could be reliably aligned, and thus were considered conserved. Examination of the evolutionary rates of the non-coding and protein-coding regions of these 732 genes identified 418 non-coding and 100 protein-coding regions whose evolutionary rate was different in *A. fumigatus* compared to close relatives. These 418 non-coding regions include 25 known genetic determinants of *A. fumigatus* virulence, such as *pkaR* (a regulatory subunit essential for protein kinase A pathway), *gliG* (glutathione S-transferase required for gliotoxin production), and *metR* (transcription factor required for sulfur assimilation); the protein-coding regions of these genes are highly conserved across species but show unique sequence differences between *A. fumigatus* and the other species in their non-coding regions. The extensive variation and different evolutionary rate of *A. fumigatus* non-coding regions may help to explain why *A. fumigatus* is a major human pathogen but most of its close relatives are not clinically relevant.

## Methods

### Genomic data collection

All *Aspergillus* genomes are publicly available and were downloaded from NCBI (https://www.ncbi.nlm.nih.gov/). These strains include *A. fumigatus* Af293 (Nierman et al., 2005), *A. fumigatus* A1163 (Fedorova et al., 2008), *A. oerlinghausenensis* CBS139183 (Steenwyk et al., 2020), *A. fischeri* NRRL1881 (Fedorova et al., 2008), *A. lentulus* IFM54703 (Kusuya et al., 2016), *A. novofumigatus* IBT 16806 (GenBank accession: MSZS00000000.1) *A. fumigatiaffinis* 5878 (Santos et al., 2020), *A. udagawae* IFM 46973 (Kusuya et al., 2016), *A. turcosus* HMR AF 1038 (Parent-Michaud et al., 2019), and *A. thermomutatus* HMR AF 39 (Parent-Michaud et al., 2019).

### Identification of single-copy orthologous genes

To infer single-copy orthologous genes among all protein-coding sequences for all ten taxa, we used OrthoFinder, version 2.4.0 (Emms & Kelly, 2015). OrthoFinder clustered genes into orthogroups algorithm Markov clustering models from gene similarity information using the sequence search program DIAMOND version 2.0.9 (Buchfink et al., 2015) and the proteomes of the ten *Aspergillus* species as input. The key parameters used in DIAMOND were e-value = 1 x 10^-3^ with a percent identity cutoff of 30% and percent match cutoff of 70%. This approach identified 5,215 single copy orthologous genes wherein all *Aspergillus* species are represented by one sequence (**Supplementary Table 1**).

### Identification of highly conserved non-coding regions

To determine a set of highly conserved non-coding regions, we first curated intergenic sequences directly upstream of the transcription start sites of all 5,215 single-copy orthologous genes for each of the ten *Aspergillus* species/strains using a custom script (https://github.com/alecbrown24/General_Bio_Scripts) based on a previously available script (https://github.com/shenwei356/bio_scripts). Up to 500 bp intergenic sequence upstream of each gene’s transcription start site were used to generate FASTA files of non-coding regions, as well as FASTA files of single-copy orthologous protein-coding sequences using Python version 3.8.2 (https://www.python.org/). Protein-coding and non-coding region FASTA sequence files were used to generate multiple sequence alignments via MAFFT version 7.453 using default parameters (Rozewicki et al., 2019). For non-coding regions, we identified a set of highly conserved multiple sequence alignments where each non-coding sequence is at least 500 bp in length and the sequence similarity of the sequence alignment between the *A. fumigatus* Af293 sequence and those of all other species is at least 75%. Analyses were conducted using custom Python scripts that used BioPython, version 1.78 (Cock et al., 2009), and NumPy, version 1.20.3 (Harris et al., 2020), modules. We discovered that 732 of the 5,215 multiple sequence alignments of non-coding regions examined were of the same length and could be reliably aligned across all species and these were used in subsequent evolutionary comparisons.

### Phylogenetic tree inference and comparisons

To construct a phylogenomic data matrix, codon-based alignments for all 5,215 single-copy protein-coding orthologs were individually trimmed using ClipKIT, version 1.1.5 (Steenwyk et al., 2020), with the ‘gappy’ mode and the gaps parameter set to 0.7. The resulting trimmed codon-based alignments were then concatenated into a single matrix with 9,248,205 sites using the ‘create_concat’ function from PhyKIT, version 1.2.1 (Steenwyk et al., 2021). Next, the evolutionary history of the ten *Aspergillus* genomes was inferred using IQ-TREE, version 2.0.6 (Minh et al., 2020), and the “GTR+F+I+G4” model of sequence evolution, which was the best fitting one according to the Bayesian Information Criterion (Waddell and Steel, 1997; Vinet and Zhedanov, 2011). Bipartition support was assessed using ultrafast bootstrap approximations (Hoang et al., 2018). All bipartitions received full support. The inferred topology is congruent with known relationships inferred from analyses of single or a few loci as well as from genome-scale analyses (Hubka et al., 2018; Steenwyk et al., 2019; dos Santos et al., 2020).

To identify gene trees whose phylogeny was statistically different from the species phylogeny, we used the approximately unbiased test (Shimodaira, 2002). Protein-coding region and non-coding region trees were inferred using IQ-TREE, version 2.0.6 (Minh et al., 2020), with “GTR+I+G+F” as it was the best fitting substitution model (Waddell and Steel, 1997; Vinet and Zhedanov, 2011). The distributions of branch lengths for protein-coding region and non-coding region trees were determined using the “total_tree_length” function from PhyKIT version 1.2.1 (Steenwyk et al., 2021). We found that 852 of the 5,215 single copy orthologs had a protein-coding tree that was statistically different from the species tree using a conservative p-value threshold of < 0.01 (**Supplementary Table 2**) For genes whose tree topologies significantly differed from the species tree, we used the corresponding gene tree for the analyses of molecular evolutionary rates of protein-coding and non-coding regions (see next section).

### Analysis of molecular evolutionary rates of protein-coding and non-coding regions between the major pathogen *A. fumigatus* and its relatives

To determine the rate of sequence evolution in protein-coding region alignments between *A. fumigatus* and close relatives, we examined variation in the ratio of the rate of nonsynonymous (dN) to the rate of synonymous (dS) substitutions (dN/dS or ω obtained codon-based alignments from their corresponding protein sequence alignments using pal2nal, version 14 (Suyama et al., 2006). We next used the codon-based alignments to calculate ω values under two different hypotheses using the codeml module in paml, version 4.9 (Yang, 2007). For each gene tested, the null hypothesis (H_0_) was that all branches of the phylogeny exhibit the same estimated ω value. We compared H_0_ to an alternative hypothesis (H_A_) which allows for the branch leading to *A. fumigatus* to have a distinct estimated ω value from the rest of the branches. To determine whether H_A_ was significantly different from H_0_ for each of the codon-based alignments, we used the likelihood ratio test with a statistical significance threshold of α = 0.01.

To determine the rate of sequence evolution in non-coding region alignments between *A. fumigatus* and close relatives, we examined variation in the ratio of the rate of substitutions in each non-coding region (dNC) to the rate of synonymous (dS) substitutions in its corresponding protein-coding region (dNC/dS or ζ) across the phylogeny. Like the analysis of the protein coding regions, the null hypothesis (H_0_) was that all branches of the phylogeny exhibit the same estimated ζ value. We compared H_0_ to an alternative hypothesis (H_A_) which allows for the branch leading to *A. fumigatus* to have a distinct estimated ζ value from the rest of the branches. ζ values were calculated under the different hypotheses using HyPhy version 2.2.2 (Pond et al., 2004) with the “nonCodingSelection.bf” batch file as established by Oliver Fedrigo (Haygood et al., 2007; Fedrigo et al., 2011). To determine whether H_A_ was significantly different from H_0_ for each of the non-coding region alignments, we used the likelihood ratio test with a statistical significance threshold of α = 0.01.

### Functional enrichment analyses of genes with signatures of different evolutionary rates

To determine whether genes with signatures of different evolutionary rates in either their coding or non-coding regions are enriched for particular functional categories, we implemented the Gene Ontology (GO) Term Finder webtool on AspGD (Cerqueira et al., 2013) using default settings. We conducted two separate analyses. The first examined the 418 *A. fumigatus* genes that exhibited a different evolutionary rate in their non-coding regions, whereas the second examined the 100 *A. fumigatus* genes with a different evolutionary rate in their protein-coding regions. These gene sets were compared to a general background set that includes all the features/gene names in the database with at least one GO annotation for *A. fumigatus*. Both analyses used a p-value cutoff of 0.05.

### Examination and visualization of mutational signatures

To identify interesting examples of sequence variation between *A. fumigatus* and the other species for non-coding regions of genes of interest, we visualized and compared multiple sequence alignments using the MView function in EMBL-EBI (Madeira et al., 2019). Workflow of methods can be seen in **Supplementary Figure 1**.

## Results

### *Aspergillus* species exhibit extensive sequence variation in their homologous non-coding regions

To analyze the sequence diversity amongst non-coding regions in section *Fumigati*, we first identified 5,215 single-copy orthologous genes amongst ten species in the section (Figure 1A). We then computed the percent similarity between the upstream region of each *A. fumigatus* Af293 ortholog and the nine homologous regions upstream from the genes in the other nine taxa. Those individual percent similarities were then averaged to get the final percent similarity for that ortholog. Averaging the upstream region percent similarities for the 5,215 single-copy orthologous genes revealed an average similarity of ∼72%; 648 alignments exhibited <50% similarity, 3,665 between 50-75% similarity, and 902 between 75-90% similarity. Despite most sequences being dissimilar, the non-coding regions of three genes exhibited > 90% similarity. These genes are *cnaB*, whose protein product is a calcineurin regulatory subunit and whose transcript is induced by exposure to human airway epithelial cells (Juvvadi et al., 2011; Oosthuizen et al., 2011), *AFUA_6G07800*, which is predicted to be a transcription factor with unknown function (Cerqueira et al., 2013), and *AFUA_6G04530*, which is predicted to have a role in histone acetylation (Cerqueira et al., 2013).

**Figure 1.**
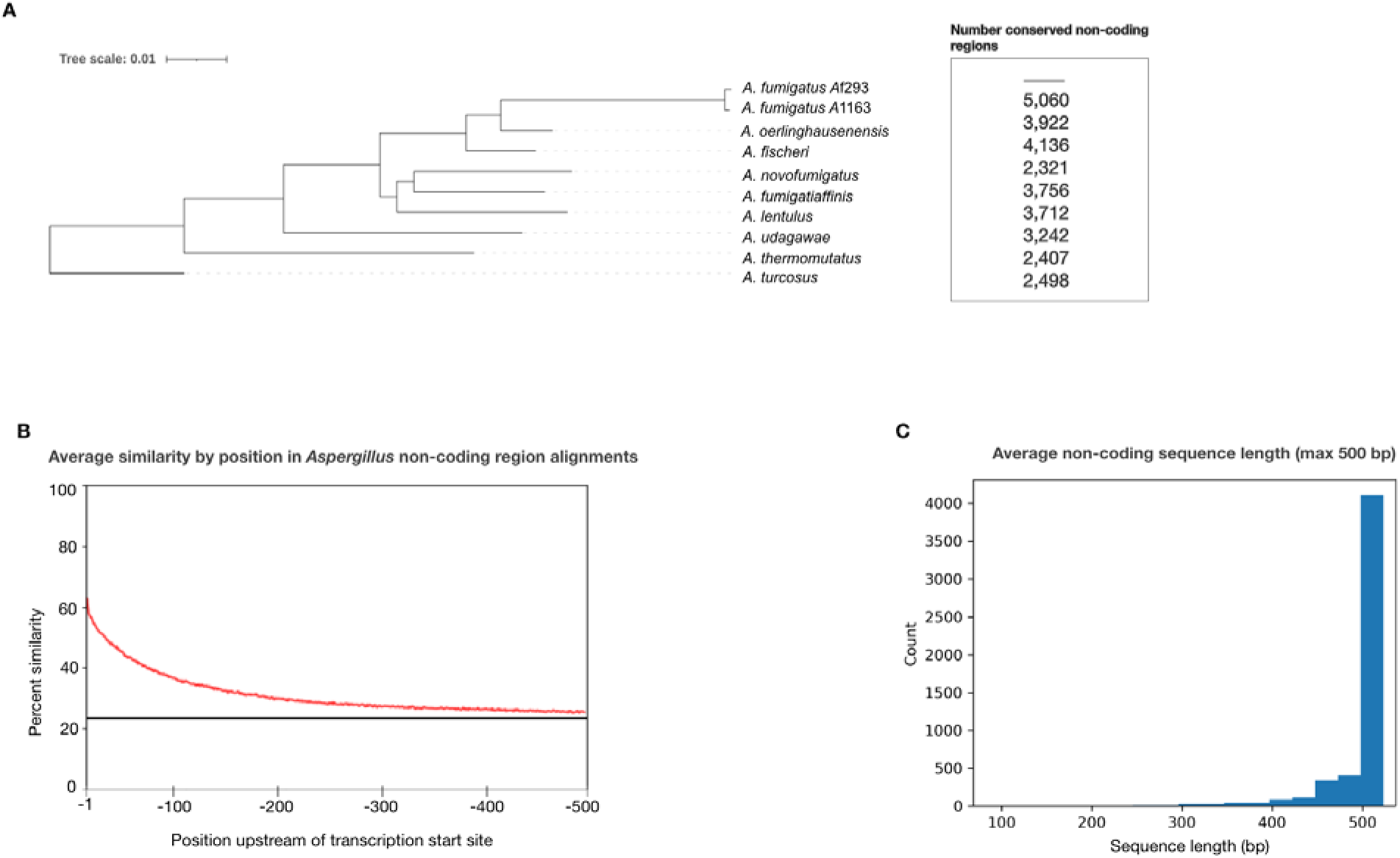
*Aspergillus* section *Fumigati* species exhibit sequence variation in non-coding regions that are 500 base pairs upstream of genes’ transcription start sites. A. Species phylogeny of two *A. fumigatus* reference strains (Af293 and A1163) and closely related *Aspergillus* section *Fumigati* species constructed from concatenation analysis of a 5,215-gene data matrix. Branch lengths correspond to nucleotide substitutions / site. Note the long branch leading to *A. fumigatus*, indicative of a greater number of nucleotide substitutions per site in this species. The number of genes whose non-coding regions were conserved (> 75% sequence similarity between each species and at least 500 bp in length) between *A. fumigatus* Af293 and each species are shown next to the corresponding taxa. B. Average percent sequence similarity of non-coding regions of 5,215 genes by position, relative to the gene’s transcription start site. Sequence alignments of non-coding regions up to 500 bp were compared by position and the average percent similarities for each site are reported with −1 indicating the site directly upstream of the transcription start site and −500 indicating the site 500 bp upstream of the transcription start site. Black horizontal line indicates 25% sequence similarity. C. Average sequence alignment lengths of the 5,215 non-coding regions examined in this study. 4,137 of the 5,215 non-coding regions have at least 500 bp in all 10 strains/species used in this study.

Average percent similarity by position in *Aspergillus* non-coding region alignments (Figure 1B) revealed that the percent similarity directly upstream of the transcription start site (TSS) (−1 bp upstream) is above 60% and decreases as the distance from the TSS increases, approaching 25% similarity – a value suggestive of lack of homology – around −500 bp upstream. This result suggests that potentially conserved promoter and *cis*-regulatory elements occur in this 500 bp region across conserved non-coding regions and is consistent with experimental data (de Castro et al., 2014; Chung et al., 2014). We also found that 4,079 regions upstream of the TSS are at least 500 bp in length across all species (Figure 1C).

Examination of the 4,567 genes whose non-coding region alignments exhibited >50% similarity revealed several instances in which one or more sequences were poorly aligned for stretches of 100 bp or more. This was especially true when sequences in these alignments exhibited large variation in their lengths. Further, we found a low level of synteny between the 5,215 non-coding regions as genes immediately upstream of a sequence of interest generally differed between species. Given the uncertainty regarding the homology of some sequences in these 5,215 non-coding region alignments and our finding that most sequence conservation was found less than 500 bp upstream of TSSs, we focused our analyses only on the 732 non-coding region alignments whose sequences were all at least 500 bp long and exhibited > 75% sequence similarity between *A. fumigatus* Af293 and each other strain / species in the phylogeny.

We also determined the number of conserved non-coding regions (>75% sequence similarity and 500 bp in length) between *A. fumigatus* Af293 and each of the other nine strains / species (Figure 1A). Interestingly, we found that *A. novofumigatus* shares the fewest number of conserved non-coding regions (2,321) despite being more closely related to *A. fumigatus* than other members of the phylogeny (Hourbraken et al., 2014; Steenwyk et al., 2019; Rokas et al., 2020); this suggests that the quality of annotations may differ across the ten genomes examined and that improvements in the gene annotation of these genomes could increase the number of conserved non-coding regions shared by these taxa. *A. fumigatus* A1163 shared the greatest number of conserved non-coding regions with 5,020 and with the exceptions of *A. novofumigatus* and *A. oerlinghausenensis*, the closer a relative is to *A. fumigatus* Af293, the greater the number of conserved non-coding regions that are shared.

### Phylogenetic analyses reveal differences between non-coding region trees and protein-coding region trees

To help determine if differences existed between non-coding and protein-coding regions across our species, we first compared total branch lengths in phylogenetic trees constructed from both non-coding and protein-coding regions from all 5,215 genes (**Supplementary Figure 2)** Comparisons of the overall distributions between these two groups revealed a statistically significant difference between the overall branch lengths of protein-coding and non-coding regions (Wilcoxon signed-ranked test; p-value = 0.004), suggesting that non-coding regions of single-copy orthologs evolve faster than protein-coding regions.

### Many non-coding but fewer protein-coding regions exhibit different rates of evolution in *A. fumigatus*

To test whether the molecular evolutionary rates of protein-coding and non-coding regions differed between the major pathogen *A. fumigatus* and its relatives, we statistically examined whether protein-coding and non-coding *A. fumigatus* sequences evolved at a similar (H_0_) or different (H_A_) rate as those of other taxa (Figure 2). Examination of protein-coding regions identified 100 genes (**Supplementary Table 3**) (13.7% of examined genes) that significantly rejected H_0_ (under which all branches exhibited the same ω value) over H_A_ (which postulates that the ω value of the branch leading to *A. fumigatus* was distinct from the background ω value of all other branches) (Figure 3B). Examination of non-coding regions identified 418 genes (**Supplementary Table 4**) (57.1% of examined genes) that significantly rejected H_0_ (under which all branches exhibited the same ζ value) over H_A_ (which postulates that the ζ value of the branch leading to *A. fumigatus* was distinct from the background ζ value of all other branches) (Figure 3C). Taken together, these results suggest a considerable amount of variation in non-coding regions than in protein-coding regions between *A. fumigatus* and relatives. Examination of the p-value distribution of protein-coding regions exhibits a uniform distribution, while the p-value distribution of non-coding regions exhibits a bimodal shape where nearly all p-values are either under 0.05 or 1.0. This result suggests that the 418 non-coding regions exhibited major differences in their relative fit for the two hypotheses, whereas protein-coding regions exhibited much smaller differences.

**Figure 2.**
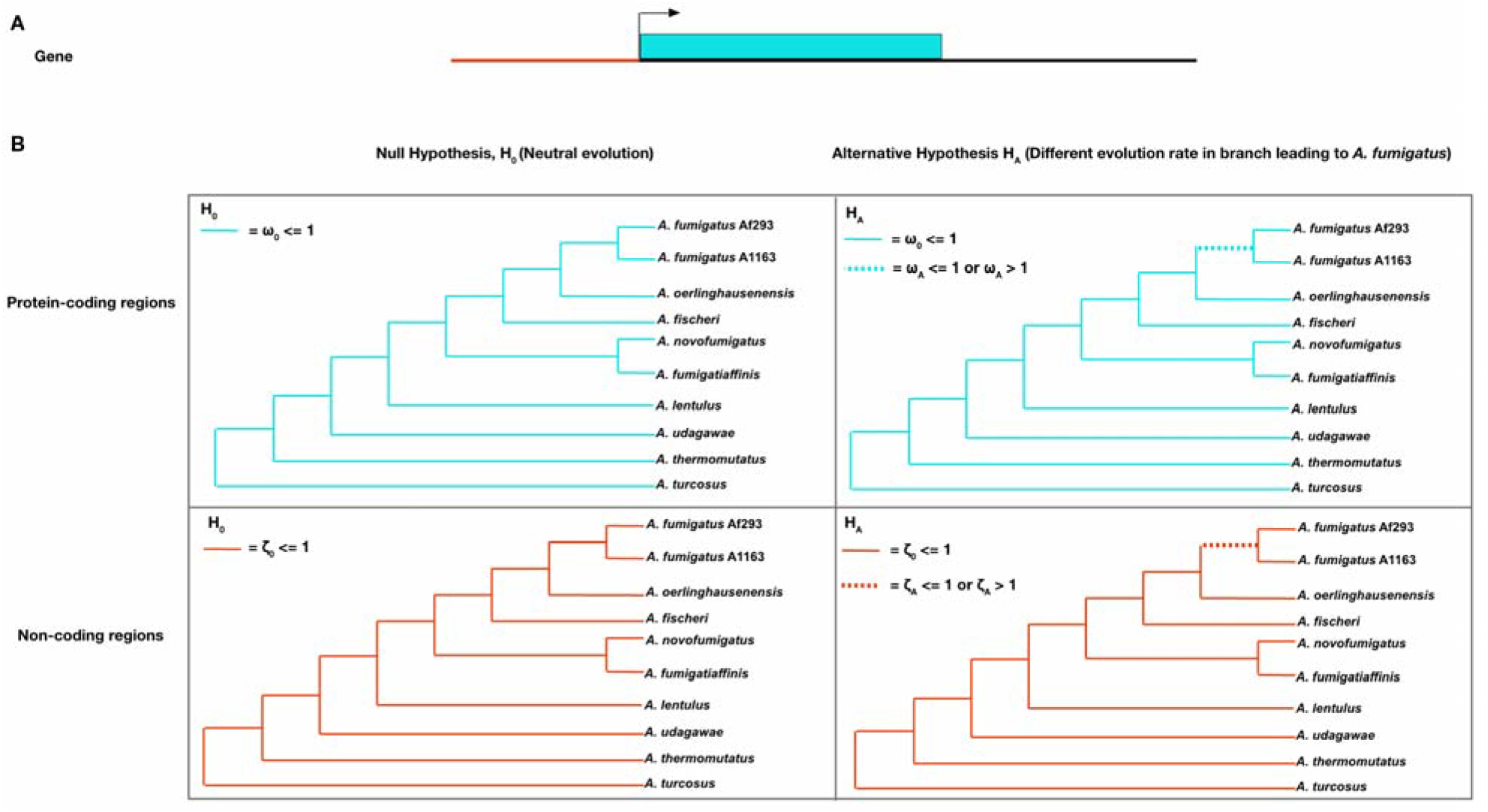
Examining whether the non-coding and protein-coding regions of 732 genes have different evolutionary rates in the major pathogen *A. fumigatus*. A. Diagram of a typical gene. The arrow represents the transcription start site; the blue box indicates the gene’s protein-coding region. The red line represents the non-coding region upstream of the gene’s transcription start site. B. The top two panels present the null and alternative hypotheses for evolutionary rate difference in the protein-coding regions of *A. fumigatus* genes relative to the other species. The null hypothesis (H_0_, upper left) constrains all ω (dN/dS) values across all branches to be less than or equal to 1, the neutral evolutionary rate. The alternative hypothesis (H_A_, upper right) allows the branch leading to *A. fumigatus* (dashed branch) to have an ω value lower than, equal to, or greater than 1 (indicative of evolutionary rate difference) compared to the background branches. The bottom two panels present the null and alternative hypotheses for evolutionary rate difference in the non-coding regions of *A. fumigatus* genes relative to other species. Similarly, the null hypothesis (H_0_, bottom left) constrains all ζ (dNC/dS) values across all branches to less than or equal to 1. The alternative hypothesis (H_A_, bottom right) allows for the branch leading to *A. fumigatus* (dashed branch) to have a ζ value lower than, equal to, or greater than 1 (evolutionary rate difference) compared to the background branches. For each protein-coding and non-coding region, a likelihood ratio test was used to determine which hypothesis best fits the data.

**Figure 3.**
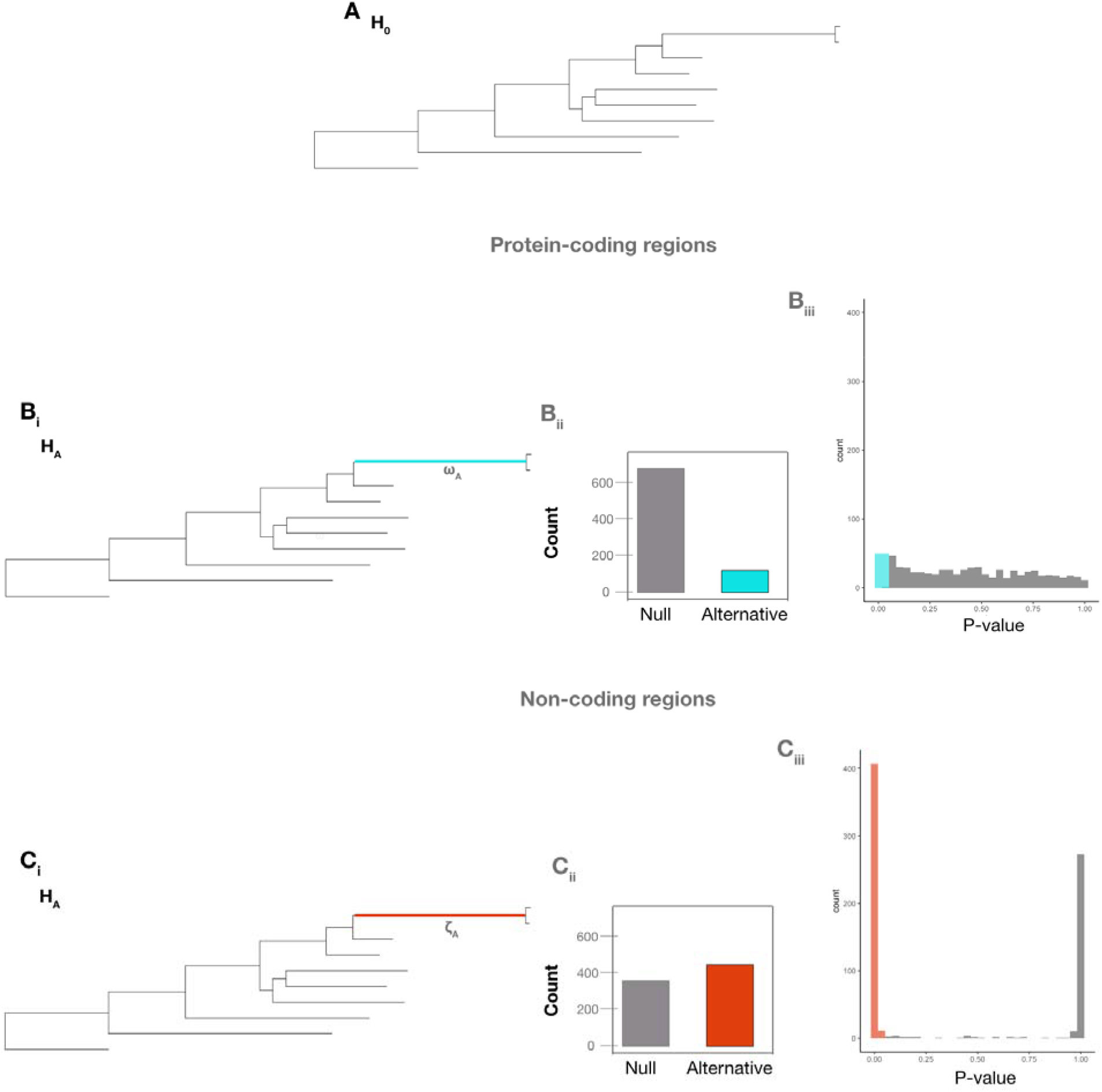
Non-coding regions of *A. fumigatus* genes exhibit many more signatures of evolutionary rate difference than their corresponding protein-coding regions. A. The null hypothesis (H_0_) that all branches have the same evolutionary rate. B-C. The alternative hypotheses assume that the ω value (Bi) or the ζ value (Ci) in the branch leading to *A. fumigatus* differs from the value in the rest of the branches of the phylogeny. Bii. 632 of 732 protein-coding regions (84.34%) did not reject H_0_ (gray) and 100 of 732 (16.66%) rejected H_0_ (blue). Biii. The distribution of p-values for protein-coding regions that did not (gray) and did (blue) reject H_0_. Cii. 314 of 732 non-coding regions (42.90%) did not reject H_0_ (gray) and 418 of 732 non-coding regions (57.10%) rejected H_0_ (red), which results suggest a greater amount of variation in non-coding regions than in protein-coding regions between *A. fumigatus* and relatives. Ciii The distribution of p-values for non-coding regions that did not (gray) and did (red) reject H_0_.

### Genes with signatures of a different evolutionary rate in non-coding regions are enriched for regulatory functions in *A. fumigatus*

To identify functions that were over-represented in the list of genes that rejected H_0_ for either their coding or non-coding regions, we conducted gene ontology (GO) enrichment analyses. Examination of significantly over-represented GO terms for the 418 genes (**Supplementary Table 5**) with signatures of different evolutionary rates in non-coding regions revealed numerous biological processes related to regulation, metabolism, and development (e.g., “cellular component organization or biogenesis”, p = 0.00016; “regulation of protein metabolic process”, p = 0.00101; “hyphal growth”, p = 0.00352; “regulation of cell cycle”; p = 0.0076; “developmental process”, p = 0.01233; “reproduction”. p = 0.00361). Of the 418 genes, 71 lacked any functional GO annotation. In comparison, for the 314 genes that did not exhibit a different evolutionary rate in their non-coding regions (**Supplementary Table 6**), the only term that was enriched was “nucleotide binding” (p = 0.06129) and was found associated with 48 genes. For the 100 genes with signatures of different evolutionary rate in their coding regions, only one function was found enriched (“regulation of cellular process”, p = 0.02647). Of note, 74 of the protein coding genes lacked any functional GO annotation (**Supplementary Table 7**).

### Four genes whose non-coding regions exhibit different evolutionary rates in *A. fumigatus* also bind transcription factors that are known genetic determinants of virulence

To identify if any of the *A. fumigatus* genes with different evolutionary rates in their respective non-coding regions also contain known TF binding sites, we compared the list of 418 non-coding regions to previously found ChIP binding sites of two TFs known to be genetic determinants of virulence, CrzA and SrbA (Cramer et al., 2008; Willger et al., 2008). ChIP-seq analysis of CrzA (de Castro et al., 2014) uncovered 110 genes that are directly bound by the TF in *A. fumigatus* strain Af293. Of these, 28 were reported to exhibit CrzA binding within 500 bp of the transcription start site, and two genes, *AFUA_8G05090* (a putative MFS transporter) and *AFUA_3G09960* (Aureobasidin resistance protein), exhibited a different evolutionary rate in the non-coding regions of *A. fumigatus* strains in our analysis. ChIP-seq analysis of SrbA (Chung et al., 2014) revealed 112 genes directly bound by the TF in *A. fumigatus* strain A1163. Of these, 57 were reported to exhibit SrbA binding within 500 bp of the transcription start site, and two genes, *AFUB_074100* (a gene of unknown function which appears to interact with *sldA*; a checkpoint protein kinase) and *AFUB_012300* (a gene predicted to be involved in nitrate assimilation), exhibited a different evolutionary rate in the non-coding regions of *A. fumigatus* strains in our analysis. Importantly, we found that for all four genes (*AFUA_8G05090*, *AFUA_3G09960*, *AFUB_074100*, and *AFUB_012300*) there was at least a 2 bp difference between the *A. fumigatus* TF binding site location and the TF binding site in at least one close relative (Figure 4). Together, our results suggest that intergenic non-coding regions that bind known TFs can exhibit substantial differences in their evolutionary rates between *A. fumigatus* and close relatives, which raises the hypothesis that these differences may lead to differences in gene expression.

**Figure 4.**
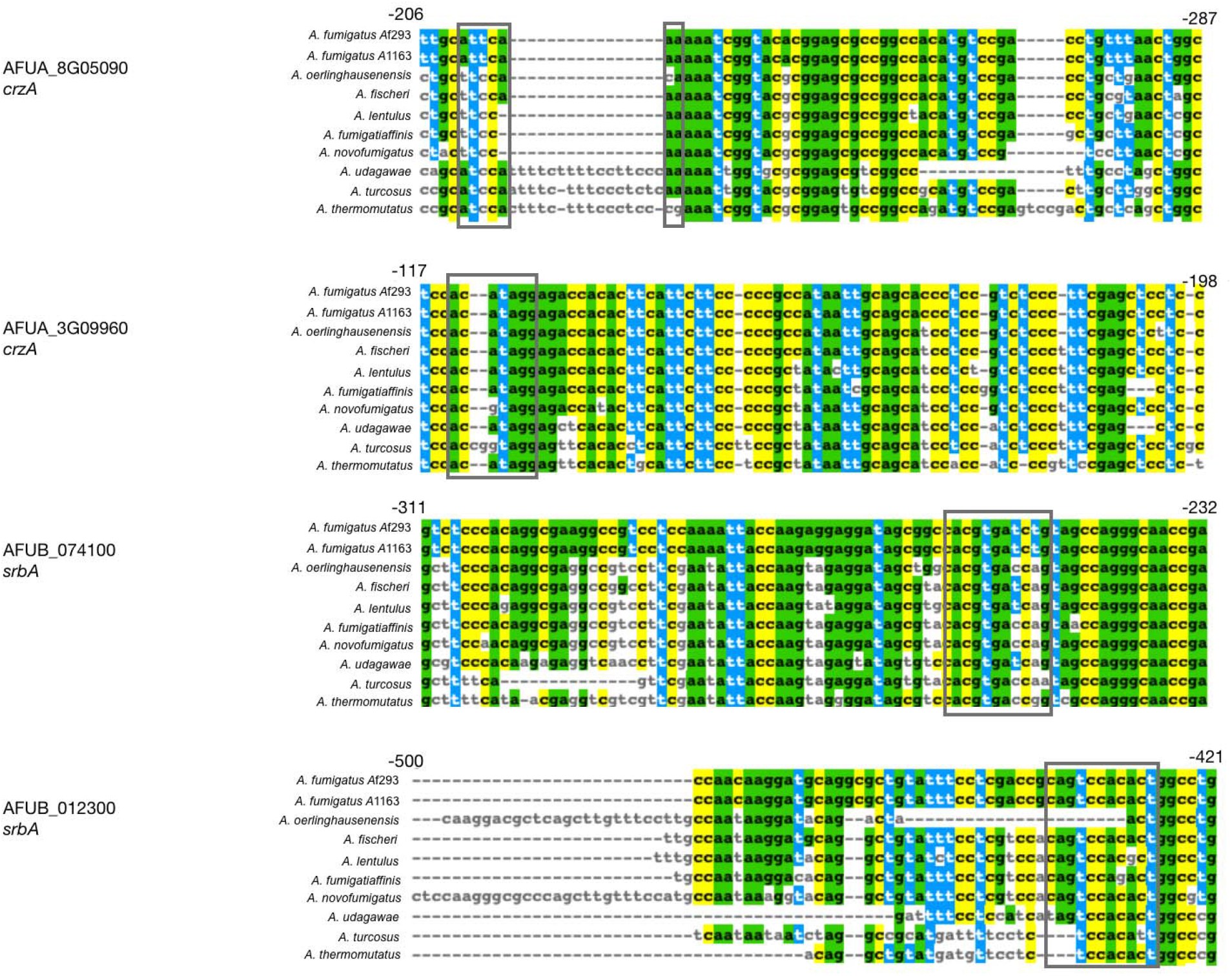
CrzA and SrbA binding locations in 4 genes that exhibit a different evolutionary rate in non-coding region. Two *A. fumigatus* genes known to bind CrzA in their non-coding region (*AFUA_8G05090* and *AFUA_3G09960*) and two *A. fumigatus* genes known to bind SrbA in their non-coding region (*AFUB_074100* and *AFUB_012300*) have at least a 2 bp difference between the *A. fumigatus* TF binding site location and the TF binding site in at least one close relative. Grey boxes represent the binding site locations found in previous ChIP-seq experiments.

### Alignments of non-coding regions upstream of genetic determinants of virulence with different rates of evolution in *A. fumigatus* reveal variants that are exclusively present or absent in *A. fumigatus*

We identified 25 genetic determinants of virulence with an upstream, non-coding region that exhibited a different rate of evolution in *A. fumigatus* (Table 1). Three genes (*metR, his3*, and *met16*) are involved in amino acid biosynthesis, eight genes (*chsF, calA, gel2, nrps1, gfa1, csmB, rlmA*, and *rodA*) are involved in cell wall biosynthesis, nine genes (*noc3, spe2, gus1, pri1, AFUA_2G10600*, *mak5, pkaR, ramA*, and *somA*) are involved in cellular metabolism, two genes (*aspB* and *tom40*) are involved in hyphal growth, and three genes (*gliG, gliI*, and *gliJ*) are involved in gliotoxin biosynthesis. Of the 25, 14 genes (*metR, chsF, calA, gel2, nrps1, gfa1, csmB, rlmA, rodA, AFUA_2G10600*, *pkaR, ramA, somA*, and *aspB*) have been shown to modulate virulence in an animal model of infectious disease. Three genes (*gliG, gliI*, and *gliJ*) are required for the biosynthesis of gliotoxin, a secondary metabolite involved in *A. fumigatus* virulence (Brakhage & Langfelder, 2002; Dagenais & Keller, 2009), and deletions of the eight remaining genes (*his3, met16, noc3, spe2, gus1, pri1, mak5*, and *tom40*) have been previously shown to be important for viability and therefore, likely virulence (for a detailed list, see: Steenwyk et al., 2021). Interestingly, only two of these 25 genes (*csmB* and *rodA*) exhibit a signature of different evolutionary rate in their protein-coding region as well, suggesting that there are more changes in homologous non-coding regions than in protein-coding regions.

**Table 1.**
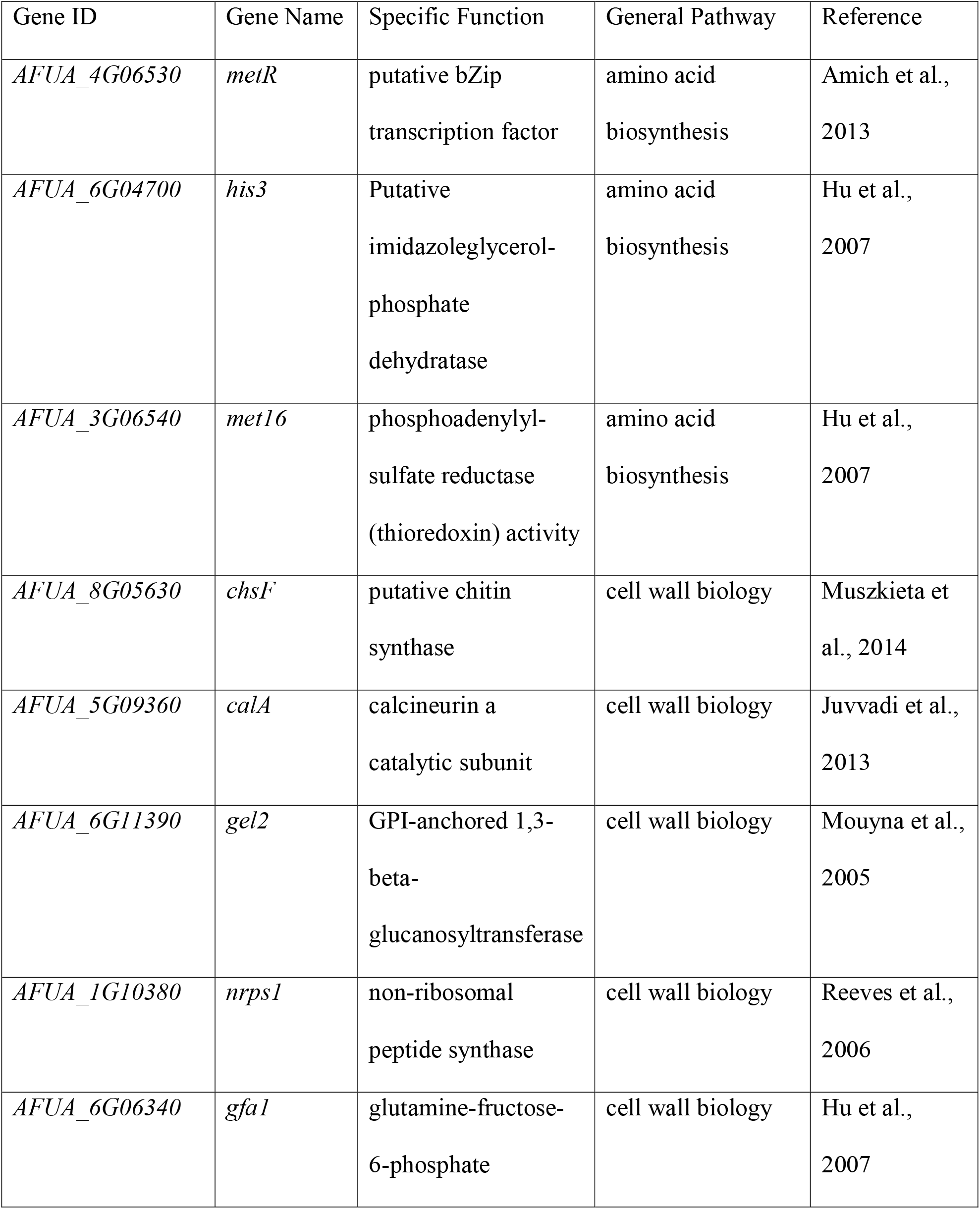
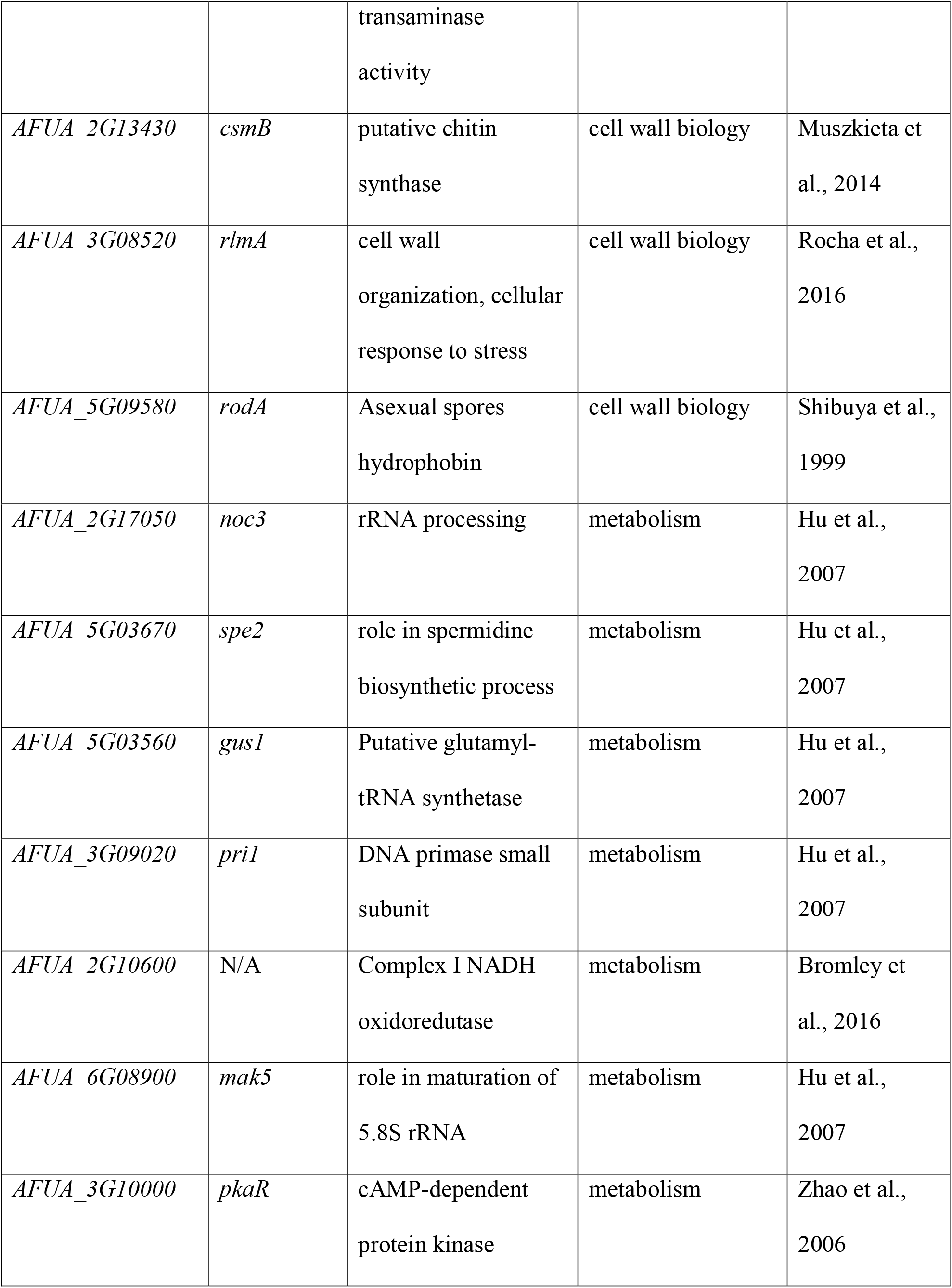
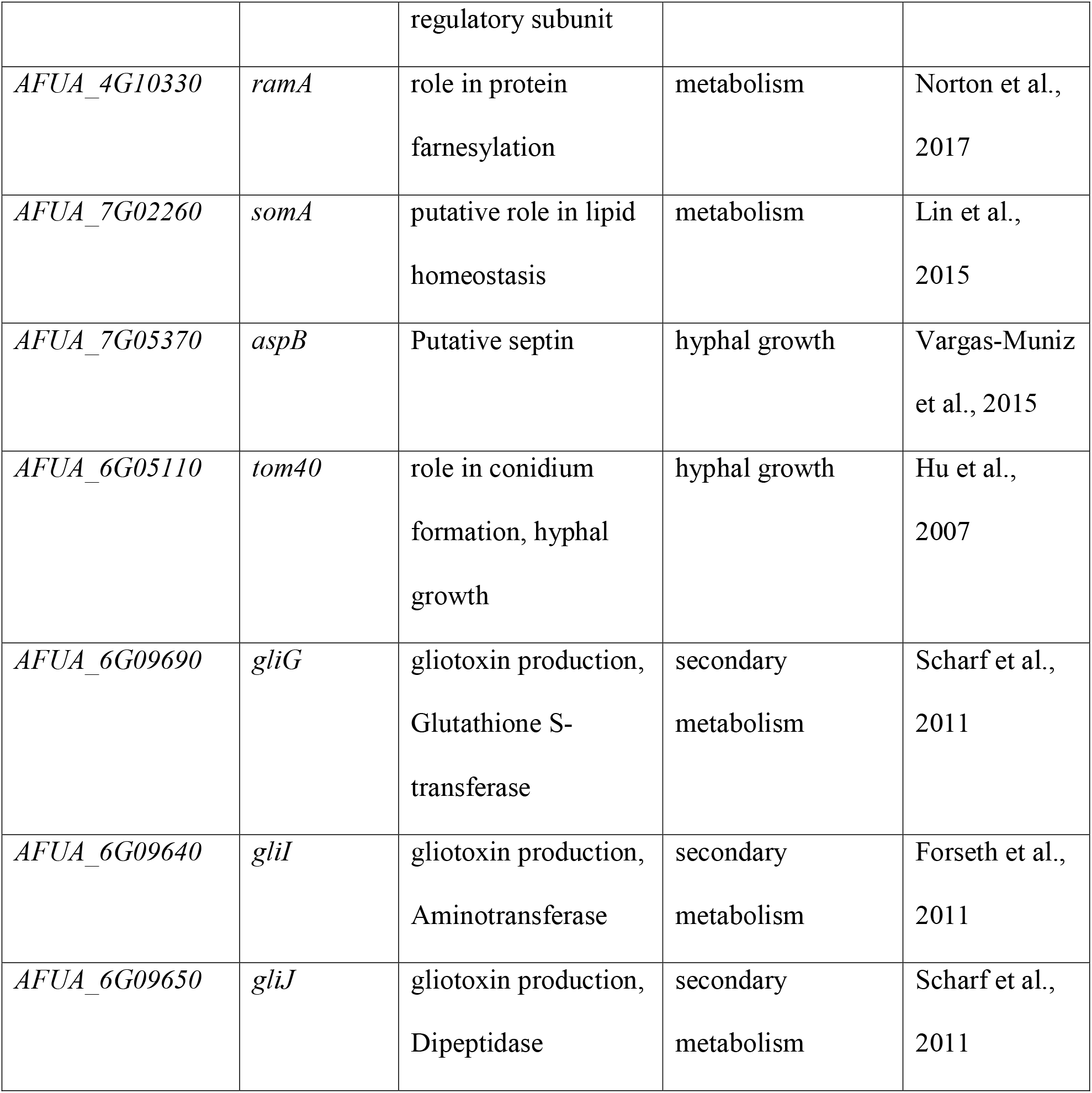
Twenty-five genetic determinants of *A. fumigatus* virulence have a different evolutionary rate in their non-coding regions.

Examination of sequence alignments of non-coding regions of these 25 genes (Table 1) reveal several interesting patterns (Figure 5). For example, the upstream non-coding region of *pkaR* exhibits a 10 bp stretch from −434 bp to −424 bp upstream of the transcription start site which is deleted exclusively in *A. fumigatus* and present and largely conserved in all other species (Figure 5A). The upstream region of g*fa1* also has a stretch of 5 bp exclusively deleted in *A. fumigatus* and present in all other species. Sequence alignment of *gliG* non-coding regions reveal an 11 bp G-rich insertion that is unique to the two *A. fumigatus* strains (Figure 5B). In addition to *A. fumigatus*-specific indels, we also observed that the non-coding regions of several genes known to be involved in *A. fumigatus* virulence exhibit indel variation across the other *Aspergillus* species examined as well. For example, the non-coding region of *metR* exhibits a 7bp pyrimidine rich insertion that is found only in *A. fumigatus*, *A. oerlinghausenensis* and *A. lentulus* (Figure 5C); while the upstream regions of *calA* and *pri1* both have small sequences (12 bp and 5 bp respectively) that are exclusively absent in *A. fumigatus* strain Af293, *A. fumigatus* strain A1163, and *A. oerlinghausenensis*.

**Figure 5.**
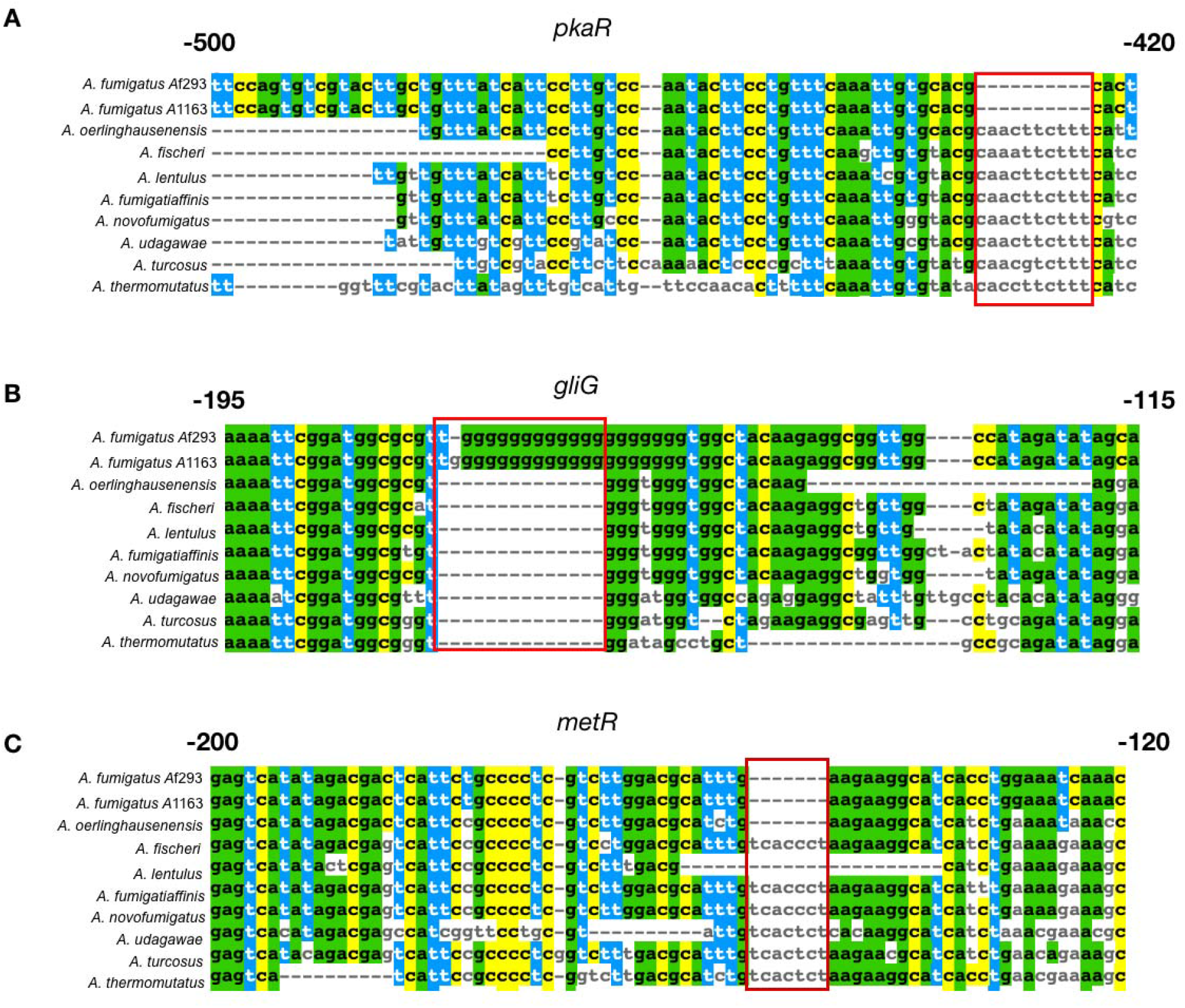
Notable examples of sequence differences between the major pathogen *A. fumigatus* and its close relatives in non-coding regions located upstream of three known genetic determinants of *A. fumigatus* virulence that exhibited signatures of a different evolutionary rate. A. *pkaR* encodes a regulatory subunit involved in the regulation of the cyclic AMP – dependent protein kinase pathway; deletion of *pkaR* in *A. fumigatus* has been shown to attenuate virulence in a neutropenic mouse model (Zhao et al., 2006). The sequence alignment of the non-coding region of the *pkaR* gene exhibits a 10 bp region (red box) that is uniquely deleted in *A. fumigatus* but present in all other close relatives. B. *gliG* encodes for a glutathione S-transferase (GST) that is part of the gliotoxin biosynthetic gene cluster and is required for gliotoxin production (Scharf et al., 2011). Gliotoxin contributes to the virulence of *A. fumigatus*, inactivating host vital proteins via conjugation (Scharf et al., 2011). The sequence alignment of the non-coding region upstream of the *gliG* gene exhibits a 15 bp G-rich region that has been inserted in *A. fumigatus* and is absent from all other species. C. *metR* encodes for a TF involved in sulfur assimilation and is a known *A. fumigatus* virulence factor (Amich et al., 2013). The sequence alignment of the non-coding region of the *metR* gene exhibits a 7 bp region that is absent in only *A. fumigatus, A. oerlinghausenensis*, and *A. lentulus*; and present in all other species. Colored nucleotides represent sites that are present in *A. fumigatus* Af293 and shared between other species, highlighting differences between the reference strain and close relatives.

## Discussion

We identified a set of 732 genes whose non-coding regions were conserved between *A. fumigatus* and close relatives and tested whether the branch leading to *A. fumigatus* exhibited a difference in the evolutionary rate in either its protein-coding or its non-coding regions compared to the other species. We found that the non-coding regions of 418 of these genes exhibit signatures of a different evolutionary rate in *A. fumigatus*. These 418 genes include 25 that are known genetic determinants of *A. fumigatus* virulence (Steenwyk et al., 2021) (Table 1). Given the differences in reported invasive aspergillosis cases caused by *A. fumigatus* compared to other *Aspergillus* species (Steinbach et al., 2012), differences in non-coding regions upstream of these 418 genes, and especially of these 25 genes previously connected to virulence, may play a role in varying pathogenic potentials of *Aspergillus* section *Fumigati* species.

Gene Ontology (GO) analysis of the 418 genes which exhibit signatures of a different evolutionary rate in non-coding regions revealed an enrichment for genes involved in regulation of metabolism and development. This is consistent with previous studies of the evolution of non-coding regions in humans (Haygood et al., 2007), which experienced positive selection in non-coding regions for genes involved in metabolism regulation (particularly glucose metabolism) and regulation of development (particularly the nervous system) compared to close relatives. These results raise the possibility that non-coding regions associated with particular functions in diverse taxa are more likely to experience changes in their evolutionary rates.

We compared our list of genes with signatures of different evolutionary rates with previous ChIP-seq studies of the transcription factors CrzA and SrbA, both of which are well studied genetic determinants of virulence for *A. fumigatus* (Cramer et al., 2008; de Castro et al., 2014; Colabardini et al., 2021; Willger et al., 2008; Chung et al., 2014). We found that the non-coding regions of two genes bound by CrzA (*AFUA_8G05090* and *AFUA_3G09960*) and two genes bound by SrbA (*AFUB_074100* and *AFUB_012300*) also exhibited different evolutionary rates in *A. fumigatus*. Interestingly, when we examined the sequence alignments of the non-coding regions of these genes, we found differences at the TF binding sites (TFBS) between *A. fumigatus* and relatives. We hypothesize that the different evolutionary rate we observed in these *A. fumigatus* genes is due to changes in the associated TFBS, which may influence the regulation of these genes.

Comparisons between *Saccharomyces cerevisiae* and *Saccharomyces paradoxus* non-coding regions that were similar to the ones we report here, revealed that TFBS tend be more conserved in the proximal promoter region (within 200bp of the transcription start site) than the distal region, yet some differences in TFBS were reported between the two species in their respective proximal promoter regions (Schaefke et al., 2015). Evolutionary differences around the TSS have also been reported in *Drosophila* species, with certain species (such as *Drosophila pseudoobscura*) exhibiting an increased mutation rate upstream of the TSS when compared to *Drosophila melanogaster* (Main et al., 2013). Combined with the results presented here, it is likely that the evolution of non-coding regions is not uniform across closely related species and that these differences may play a functional role in downstream gene expression.

We compared our list of 418 genes with signatures of a different evolutionary rate in the non-coding regions of *A. fumigatus* to a previously curated set of 206 genetic determinants of *A. fumigatus* virulence (Steenwyk et al., 2021) and found that 25 genes were shared between the two datasets. We also found that the most represented general function amongst these 25 genes was “metabolism”, which raises the question of their impact on virulence, given the role that metabolism has been shown to play in *A. fumigatus* virulence (Willger et al., 2009). In particular, *pkaR* is essential for proper protein kinase A signaling (Griffioen & Thevelein, 2002) and plays a key role in the germination and growth of *A. fumigatus* asexual spores (Zhao et al., 2006). Moreover, *pkaR* has been shown to be required for *A. fumigatus* virulence (Lin et al., 2015) in an immunocompromised murine model of invasive aspergillosis (Fuller et al., 2011). We found that the protein-coding region of *pkaR* does not exhibit a different evolutionary rate in *A. fumigatus*, suggesting that the gene’s protein-coding region is conserved. However, analysis of the sequence alignment of the non-coding region revealed a 11 bp region (CAACTTCTTT) absent in *A. fumigatus* but present in all other species (Figure 5). Interestingly, this binding site is similar to the predicted TFBS for the *S. cerevisiae* TF Ste12 (Badis et al., 2008), a homolog to SteA in *A. fumigatus*. While it has yet to be elucidated if this particular site upstream of *pkaR* is involved in SteA binding, it may be that the absence of this specific 11 bp region may change the regulation of *pkaR* and thus somehow contribute to *A. fumigatus* virulence.

The role of gliotoxin in *A. fumigatus*-mediated disease has been of increasing interest, due to its ability to inhibit the host immune response (Raffa & Keller, 2019). However, the gliotoxin biosynthetic gene cluster is found in both *A. fumigatus* and its non-pathogenic close relatives *A. oerlinghausenensis* and *A. fischeri*, and all three species are known to produce gliotoxin (Knowles et al., 2020; Steenwyk et al., 2020). Here, we identify that three genes in the gliotoxin biosynthetic gene cluster (*gliG, gliJ, gliI*) exhibit a different evolutionary rate in their upstream non-coding regions in *A. fumigatus*. Gliotoxin genes have been shown to require certain TFs (GliZ and RglT for example) for gliotoxin biosynthesis and/or self-protection (Schrettl et al., 2010; Ries et al., 2020; de Castro et al., 2021). Interestingly, analysis of the sequence alignment of the non-coding region of *gliG* revealed a G-rich region unique to *A. fumigatus* (Figure 5). G-rich regions have been previously reported to be found in biologically active sites and to play important roles in regulating cellular processes such as gene expression (Maizels & Gray, 2013; Maity et al., 2020). This *A. fumigatus*-specific G-rich region may contribute to some unknown gliotoxin expression pattern that contributes to *A. fumigatus* virulence or the lack of disease caused by other closely related *Aspergillus* species.

*metR* encodes a bZIP DNA binding protein required for sulfur metabolism in *A. fumigatus* and whose gene expression is regulated by LaeA, a major regulator of secondary metabolism (Jain et al., 2018. Pertaining to *A. fumigatus* virulence, sulfur assimilation plays key roles in oxidative stress response and gliotoxin biosynthesis (Traynor et al., 2019). Recent efforts have identified differences in the transcriptional profiles of *A. fumigatus* and relatives in response to exogenous gliotoxin, highlighting the pathways relating sulfur assimilation and gliotoxin production (de Castro et al., 2021). The non-coding region of *metR* possesses a 7 bp region (TCACCT) in *A. fischeri* and five other species; in contrast, the two strains of *A. fumigatus*, *A. oerlinghausenensis* and *A. lentulu*s all lack this 7 bp motif (Figure 5). While it remains unclear if this 7 bp motif has a functional role in the expression of *metR*, this result nicely illustrates the complex patterns of sequence evolution of non-coding regions in this clade of pathogens and non-pathogens.

Future work will functionally test if the non-coding region differences we report here play a role in *A. fumigatus* expression and virulence. Further, testing if non-coding regions in a larger set of *A. fumigatus* strains exhibit differences in evolutionary rates would help to elucidate more recent evolutionary changes in *A. fumigatus* and the pathogenic differences observed in these strains as well.

## Acknowledgements

We thank the Rokas lab for helpful discussion and feedback. A.B. was funded by the Biological Sciences graduate program at Vanderbilt University. Research in A.R.’s lab is supported by grants from the National Science Foundation (DEB-1442113 and DEB-2110404), the National Institutes of Health/National Institute of Allergy and Infectious Diseases (R56 AI146096), and the Burroughs Wellcome Fund. J.L.S. and A.R. were supported by the Howard Hughes Medical Institute through the James H. Gilliam Fellowships for Advanced Study program.

## References

Amich, J., Schafferer, L., Haas, H., and Krappmann, S. (2013). Regulation of Sulphur Assimilation Is Essential for Virulence and Affects Iron Homeostasis of the Human-Pathogenic Mould Aspergillus fumigatus. PLOS Pathogens 9, e1003573. doi:10.1371/journal.ppat.1003573.

Badis, G., Chan, E. T., van Bakel, H., Pena-Castillo, L., Tillo, D., Tsui, K., et al. (2008). A library of yeast transcription factor motifs reveals a widespread function for Rsc3 in targeting nucleosome exclusion at promoters. Mol Cell 32, 878–887. doi:10.1016/j.molcel.2008.11.020.

Bertuzzi, M., Hayes, G. E., Icheoku, U. J., van Rhijn, N., Denning, D. W., Osherov, N., et al. (2018). Anti-Aspergillus Activities of the Respiratory Epithelium in Health and Disease. J Fungi (Basel) 4, E8. doi:10.3390/jof4010008.

Bongomin, F., Gago, S., Oladele, R. O., and Denning, D. W. (2017). Global and Multi-National Prevalence of Fungal Diseases—Estimate Precision. J Fungi (Basel) 3, 57. doi:10.3390/jof3040057.

Brakhage, A. A., and Langfelder, K. (2002). Menacing mold: the molecular biology of Aspergillus fumigatus. Annu Rev Microbiol 56, 433–455. doi:10.1146/annurev.micro.56.012302.160625.

Bromley, M., Johns, A., Davies, E., Fraczek, M., Gilsenan, J. M., Kurbatova, N., et al. (2016). Mitochondrial Complex I Is a Global Regulator of Secondary Metabolism, Virulence and Azole Sensitivity in Fungi. PLOS ONE 11, e0158724. doi:10.1371/journal.pone.0158724.

Buchfink, B., Xie, C., and Huson, D. H. (2015). Fast and sensitive protein alignment using DIAMOND. Nat Methods 12, 59–60. doi:10.1038/nmeth.3176.

Caron, B., Luo, Y., and Rausell, A. (2019). NCBoost classifies pathogenic non-coding variants in Mendelian diseases through supervised learning on purifying selection signals in humans. Genome Biol 20, 32. doi:10.1186/s13059-019-1634-2.

Carroll, S. B. (2008). Evo-Devo and an Expanding Evolutionary Synthesis: A Genetic Theory of Morphological Evolution. Cell 134, 25–36. doi:10.1016/j.cell.2008.06.030.

Castro, P. A. de, Moraes, M., Colabardini, A. C., Horta, M. A. C., Knowles, S. L., Raja, H. A., et al. (2021). Regulation of gliotoxin biosynthesis and protection in Aspergillus species. doi:10.1101/2021.08.16.456458.

Cerqueira, G. C., Arnaud, M. B., Inglis, D. O., Skrzypek, M. S., Binkley, G., Simison, M., et al. (2014). The Aspergillus Genome Database: multispecies curation and incorporation of RNA-Seq data to improve structural gene annotations. Nucleic Acids Res 42, D705–710. doi:10.1093/nar/gkt1029.

Chakrabarti, A., Kaur, H., Savio, J., Rudramurthy, S. M., Patel, A., Shastri, P., et al. (2019). Epidemiology and clinical outcomes of invasive mould infections in Indian intensive care units (FISF study). Journal of Critical Care 51, 64–70. doi:10.1016/j.jcrc.2019.02.005.

Chotirmall, S. H., Al-Alawi, M., Mirkovic, B., Lavelle, G., Logan, P. M., Greene, C. M., et al. (2013). Aspergillus-Associated Airway Disease, Inflammation, and the Innate Immune Response. Biomed Res Int 2013, 723129. doi:10.1155/2013/723129.

Chung, D., Barker, B. M., Carey, C. C., Merriman, B., Werner, E. R., Lechner, B. E., et al. (2014). ChIP-seq and in vivo transcriptome analyses of the Aspergillus fumigatus SREBP SrbA reveals a new regulator of the fungal hypoxia response and virulence. PLoS Pathog 10, e1004487. doi:10.1371/journal.ppat.1004487.

Cock, P. J. A., Antao, T., Chang, J. T., Chapman, B. A., Cox, C. J., Dalke, A., et al. (2009). Biopython: freely available Python tools for computational molecular biology and bioinformatics. Bioinformatics 25, 1422–1423. doi:10.1093/bioinformatics/btp163.

Colabardini, A. C., Wang, F., Dong, Z., Pardeshi, L., Rocha, M. C., Costa, J. H., et al. (2021). Heterogeneity in the transcriptional response of the human pathogen Aspergillus fumigatus to the antifungal agent caspofungin. doi:10.1101/2021.07.15.452449.

Cramer, R. A., Perfect, B. Z., Pinchai, N., Park, S., Perlin, D. S., Asfaw, Y. G., et al. (2008). Calcineurin target CrzA regulates conidial germination, hyphal growth, and pathogenesis of Aspergillus fumigatus. Eukaryot Cell 7, 1085–1097. doi:10.1128/EC.00086-08.

Croft, C. A., Culibrk, L., Moore, M. M., and Tebbutt, S. J. (2016). Interactions of Aspergillus fumigatus Conidia with Airway Epithelial Cells: A Critical Review. Front Microbiol 7, 472. doi:10.3389/fmicb.2016.00472.

Dagenais, T. R. T., and Keller, N. P. (2009). Pathogenesis of Aspergillus fumigatus in Invasive Aspergillosis. Clin Microbiol Rev 22, 447–465. doi:10.1128/CMR.00055-08.

de Castro, P. A., Chen, C., de Almeida, R. S. C., Freitas, F. Z., Bertolini, M. C., Morais, E. R., et al. (2014). ChIP-seq reveals a role for CrzA in the Aspergillus fumigatus high-osmolarity glycerol response (HOG) signalling pathway. Mol Microbiol 94, 655–674. doi:10.1111/mmi.12785.

de Vries, R. P., Riley, R., Wiebenga, A., Aguilar-Osorio, G., Amillis, S., Uchima, C. A., et al. (2017). Comparative genomics reveals high biological diversity and specific adaptations in the industrially and medically important fungal genus Aspergillus. Genome Biol 18, 28. doi:10.1186/s13059-017-1151-0.

Eide, D. J. (2020). Transcription factors and transporters in zinc homeostasis: lessons learned from fungi. Crit Rev Biochem Mol Biol 55, 88–110. doi:10.1080/10409238.2020.1742092.

Emms, D. M., and Kelly, S. (2015). OrthoFinder: solving fundamental biases in whole genome comparisons dramatically improves orthogroup inference accuracy. Genome Biology 16, 157. doi:10.1186/s13059-015-0721-2.

Fedorova, N. D., Khaldi, N., Joardar, V. S., Maiti, R., Amedeo, P., Anderson, M. J., et al. (2008). Genomic islands in the pathogenic filamentous fungus Aspergillus fumigatus. PLoS Genet 4, e1000046. doi:10.1371/journal.pgen.1000046.

Fedrigo, O., Pfefferle, A. D., Babbitt, C. C., Haygood, R., Wall, C. E., and Wray, G. A. (2011). A Potential Role for Glucose Transporters in the Evolution of Human Brain Size. Brain Behav Evol 78, 315–326. doi:10.1159/000329852.

Flores, M. E. B., Medina, P. G., Camacho, S. P. D., de Jesús Uribe Beltrán, M., De la Cruz Otero, M. del C., Ramírez, I. O., et al. (2014). Fungal spore concentrations in indoor and outdoor air in university libraries, and their variations in response to changes in meteorological variables. Int J Environ Health Res 24, 320–340. doi:10.1080/09603123.2013.835029.

Forseth, R. R., Fox, E. M., Chung, D., Howlett, B. J., Keller, N. P., and Schroeder, F. C. (2011). Identification of cryptic products of the gliotoxin gene cluster using NMR-based comparative metabolomics and a model for gliotoxin biosynthesis. J Am Chem Soc 133, 9678–9681. doi:10.1021/ja2029987.

Fuller, K. K., Richie, D. L., Feng, X., Krishnan, K., Stephens, T. J., Wikenheiser-Brokamp, K. A., et al. (2011). Divergent Protein Kinase A isoforms co-ordinately regulate conidial germination, carbohydrate metabolism and virulence in Aspergillus fumigatus. Mol Microbiol 79, 1045–1062. doi:10.1111/j.1365-2958.2010.07509.x.

Furukawa, T., van Rhijn, N., Fraczek, M., Gsaller, F., Davies, E., Carr, P., et al. (2020). The negative cofactor 2 complex is a key regulator of drug resistance in Aspergillus fumigatus. Nat Commun 11, 427. doi:10.1038/s41467-019-14191-1.

Gago, S., Overton, N. L. D., Ben-Ghazzi, N., Novak-Frazer, L., Read, N. D., Denning, D. W., et al. (2018). Lung colonization by Aspergillus fumigatus is controlled by ZNF77. Nat Commun 9, 3835. doi:10.1038/s41467-018-06148-7.

Griffioen, G., and Thevelein, J. M. (2002). Molecular mechanisms controlling the localisation of protein kinase A. Curr Genet 41, 199–207. doi:10.1007/s00294-002-0308-9.

Harris, C. R., Millman, K. J., van der Walt, S. J., Gommers, R., Virtanen, P., Cournapeau, D., et al. (2020). Array programming with NumPy. Nature 585, 357–362. doi:10.1038/s41586-020-2649-2.

Haygood, R., Fedrigo, O., Hanson, B., Yokoyama, K.-D., and Wray, G. A. (2007). Promoter regions of many neural-and nutrition-related genes have experienced positive selection during human evolution. Nat Genet 39, 1140–1144. doi:10.1038/ng2104.

Hoang, D. T., Chernomor, O., von Haeseler, A., Minh, B. Q., and Vinh, L. S. (2018). UFBoot2: Improving the Ultrafast Bootstrap Approximation. Mol Biol Evol 35, 518–522. doi:10.1093/molbev/msx281.

Houbraken, J., de Vries, R. P., and Samson, R. A. (2014). “Chapter Four - Modern Taxonomy of Biotechnologically Important Aspergillus and Penicillium Species,” in Advances in Applied Microbiology, eds. S. Sariaslani and G. M. Gadd (Academic Press), 199–249. doi:10.1016/B978-0-12-800262-9.00004-4.

Houbraken, J., Kocsubé, S., Visagie, C. M., Yilmaz, N., Wang, X.-C., Meijer, M., et al. (2020). Classification of Aspergillus, Penicillium, Talaromyces and related genera (Eurotiales): An overview of families, genera, subgenera, sections, series and species. Studies in Mycology 95, 5–169. doi:10.1016/j.simyco.2020.05.002.

Hu, W., Sillaots, S., Lemieux, S., Davison, J., Kauffman, S., Breton, A., et al. (2007). Essential Gene Identification and Drug Target Prioritization in Aspergillus fumigatus. PLOS Pathogens 3, e24. doi:10.1371/journal.ppat.0030024.

Hubka, V., Barrs, V., Dudová, Z., Sklenář, F., Kubátová, A., Matsuzawa, T., et al. (2018). Unravelling species boundaries in the Aspergillus viridinutans complex (section Fumigati): opportunistic human and animal pathogens capable of interspecific hybridization. Persoonia 41, 142–174. doi:10.3767/persoonia.2018.41.08.

Jain, S., Sekonyela, R., Knox, B. P., Palmer, J. M., Huttenlocher, A., Kabbage, M., et al. (2018). Selenate sensitivity of a laeA mutant is restored by overexpression of the bZIP protein MetR in Aspergillus fumigatus. Fungal Genet Biol 117, 1–10. doi:10.1016/j.fgb.2018.05.001.

Jang, Y. J., LaBella, A. L., Feeney, T. P., Braverman, N., Tuchman, M., Morizono, H., et al. (2018). Disease-causing mutations in the promoter and enhancer of the ornithine transcarbamylase gene. Hum Mutat 39, 527–536. doi:10.1002/humu.23394.

Juvvadi, P. R., Fortwendel, J. R., Rogg, L. E., Burns, K. A., Randell, S. H., and Steinbach, W. J. (2011). Localization and activity of the calcineurin catalytic and regulatory subunit complex at the septum is essential for hyphal elongation and proper septation in Aspergillus fumigatus. Mol Microbiol 82, 1235–1259. doi:10.1111/j.1365-2958.2011.07886.x.

Juvvadi, P. R., Gehrke, C., Fortwendel, J. R., Lamoth, F., Soderblom, E. J., Cook, E. C., et al. (2013). Phosphorylation of Calcineurin at a Novel Serine-Proline Rich Region Orchestrates Hyphal Growth and Virulence in Aspergillus fumigatus. PLOS Pathogens 9, e1003564. doi:10.1371/journal.ppat.1003564.

Kim, M. S., Cho, K. H., Park, K. H., Jang, J., and Hahn, J.-S. (2019). Activation of Haa1 and War1 transcription factors by differential binding of weak acid anions in Saccharomyces cerevisiae. Nucleic Acids Res 47, 1211–1224. doi:10.1093/nar/gky1188.

Knowles, S. L., Mead, M. E., Silva, L. P., Raja, H. A., Steenwyk, J. L., Goldman, G. H., et al. (2020). Gliotoxin, a Known Virulence Factor in the Major Human Pathogen Aspergillus fumigatus, Is Also Biosynthesized by Its Nonpathogenic Relative Aspergillus fischeri. mBio 11, e03361–19. doi:10.1128/mBio.03361-19.

Kusuya, Y., Sakai, K., Kamei, K., Takahashi, H., and Yaguchi, T. (2016). Draft Genome Sequence of the Pathogenic Filamentous Fungus Aspergillus lentulus IFM 54703T. Genome Announc 4, e01568–15. doi:10.1128/genomeA.01568-15.

Li, H., and Johnson, A. D. (2010). Evolution of Transcription Networks — Lessons from Yeasts. Curr Biol 20, R746–R753. doi:10.1016/j.cub.2010.06.056.

Lin, C.-J., Sasse, C., Gerke, J., Valerius, O., Irmer, H., Frauendorf, H., et al. (2015). Transcription Factor SomA Is Required for Adhesion, Development and Virulence of the Human Pathogen Aspergillus fumigatus. PLOS Pathogens 11, e1005205. doi:10.1371/journal.ppat.1005205.

Madeira, F., Park, Y. mi, Lee, J., Buso, N., Gur, T., Madhusoodanan, N., et al. (2019). The EMBL-EBI search and sequence analysis tools APIs in 2019. Nucleic Acids Research 47, W636–W641. doi:10.1093/nar/gkz268.

Main, B. J., Smith, A. D., Jang, H., and Nuzhdin, S. V. (2013). Transcription Start Site Evolution in Drosophila. Molecular Biology and Evolution 30, 1966–1974. doi:10.1093/molbev/mst085.

Maity, A., Winnerdy, F. R., Chang, W. D., Chen, G., and Phan, A. T. (2020). Intra-locked G-quadruplex structures formed by irregular DNA G-rich motifs. Nucleic Acids Research 48, 3315–3327. doi:10.1093/nar/gkaa008.

Maizels, N., and Gray, L. T. (2013). The G4 genome. PLoS Genet 9, e1003468. doi:10.1371/journal.pgen.1003468.

Mead, M. E., Knowles, S. L., Raja, H. A., Beattie, S. R., Kowalski, C. H., Steenwyk, J. L., et al. (2019). Characterizing the Pathogenic, Genomic, and Chemical Traits of Aspergillus fischeri, a Close Relative of the Major Human Fungal Pathogen Aspergillus fumigatus. 4, 18.

Mead, M. E., Steenwyk, J. L., Silva, L. P., de Castro, P. A., Saeed, N., Hillmann, F., et al. (2021). An evolutionary genomic approach reveals both conserved and species-specific genetic elements related to human disease in closely related Aspergillus fungi. Genetics 218, iyab066. doi:10.1093/genetics/iyab066.

Minh, B. Q., Schmidt, H. A., Chernomor, O., Schrempf, D., Woodhams, M. D., von Haeseler, A., et al. (2020). IQ-TREE 2: New Models and Efficient Methods for Phylogenetic Inference in the Genomic Era. Molecular Biology and Evolution 37, 1530–1534. doi:10.1093/molbev/msaa015.

Mouyna, I., Morelle, W., Vai, M., Monod, M., Léchenne, B., Fontaine, T., et al. (2005). Deletion of GEL2 encoding for a beta(1-3)glucanosyltransferase affects morphogenesis and virulence in Aspergillus fumigatus. Mol Microbiol 56, 1675–1688. doi:10.1111/j.1365-2958.2005.04654.x.

Muszkieta, L., Aimanianda, V., Mellado, E., Gribaldo, S., Alcàzar-Fuoli, L., Szewczyk, E., et al. (2014). Deciphering the role of the chitin synthase families 1 and 2 in the in vivo and in vitro growth of Aspergillus fumigatus by multiple gene targeting deletion. Cell Microbiol 16, 1784–1805. doi:10.1111/cmi.12326.

Nierman, W. C., Pain, A., Anderson, M. J., Wortman, J. R., Kim, H. S., Arroyo, J., et al. (2005). Genomic sequence of the pathogenic and allergenic filamentous fungus Aspergillus fumigatus. Nature 438, 1151–1156. doi:10.1038/nature04332.

Norton, T. S., Al Abdallah, Q., Hill, A. M., Lovingood, R. V., and Fortwendel, J. R. (2017). The Aspergillus fumigatus farnesyltransferase β-subunit, RamA, mediates growth, virulence, and antifungal susceptibility. Virulence 8, 1401–1416. doi:10.1080/21505594.2017.1328343.

Oosthuizen, J. L., Gomez, P., Ruan, J., Hackett, T. L., Moore, M. M., Knight, D. A., et al. (2011). Dual organism transcriptomics of airway epithelial cells interacting with conidia of Aspergillus fumigatus. PLoS One 6, e20527. doi:10.1371/journal.pone.0020527.

Parent-Michaud, M., Dufresne, P. J., Fournier, É., Martineau, C., Moreira, S., Perkins, V., et al. Draft Genome Sequences of Azole-Resistant and Azole-Susceptible Aspergillus turcosus Clinical Isolates Recovered from Bronchoalveolar Lavage Fluid Samples. Microbiology Resource Announcements 8, e01446–18. doi:10.1128/MRA.01446-18.

Pond, S. L. K., Frost, S. D. W., and Muse, S. V. (2005). HyPhy: hypothesis testing using phylogenies. Bioinformatics 21, 676–679. doi:10.1093/bioinformatics/bti079.

Raffa, N., and Keller, N. P. (2019). A call to arms: Mustering secondary metabolites for success and survival of an opportunistic pathogen. PLoS Pathog 15, e1007606. doi:10.1371/journal.ppat.1007606.

Reeves, E. P., Reiber, K., Neville, C., Scheibner, O., Kavanagh, K., and Doyle, S. (2006). A nonribosomal peptide synthetase (Pes1) confers protection against oxidative stress in Aspergillus fumigatus. FEBS J 273, 3038–3053. doi:10.1111/j.1742-4658.2006.05315.x.

Ries, L. N. A., Pardeshi, L., Dong, Z., Tan, K., Steenwyk, J. L., Colabardini, A. C., et al. (2020). The Aspergillus fumigatus transcription factor RglT is important for gliotoxin biosynthesis and self-protection, and virulence. PLoS Pathog 16, e1008645. doi:10.1371/journal.ppat.1008645.

Rocha, M. C., Fabri, J. H. T. M., Franco de Godoy, K., Alves de Castro, P., Hori, J. I., Ferreira da Cunha, A., et al. (2016). Aspergillus fumigatus MADS-Box Transcription Factor rlmA Is Required for Regulation of the Cell Wall Integrity and Virulence. G3 (Bethesda) 6, 2983–3002. doi:10.1534/g3.116.031112.

Rokas, A., Mead, M. E., Steenwyk, J. L., Oberlies, N. H., and Goldman, G. H. (2020). Evolving moldy murderers: Aspergillus section Fumigati as a model for studying the repeated evolution of fungal pathogenicity. PLOS Pathogens 16, e1008315. doi:10.1371/journal.ppat.1008315.

Ropero, P., Erquiaga, S., Arrizabalaga, B., Pérez, G., de la Iglesia, S., Torrejón, M. J., et al. (2017). Phenotype of mutations in the promoter region of the β-globin gene. J Clin Pathol 70, 874–878. doi:10.1136/jclinpath-2017-204378.

Rozewicki, J., Li, S., Amada, K. M., Standley, D. M., and Katoh, K. (2019). MAFFT-DASH: integrated protein sequence and structural alignment. Nucleic Acids Research 47, W5–W10. doi:10.1093/nar/gkz342.

Santos, R. A. C. dos, Steenwyk, J. L., Rivero-Menendez, O., Mead, M. E., Silva, L. P., Bastos, R. W., et al. (2020). Genomic and phenotypic heterogeneity of clinical isolates of the human pathogens Aspergillus fumigatus, Aspergillus lentulus and Aspergillus fumigatiaffinis. bioRxiv, 2020.02.28.970384. doi:10.1101/2020.02.28.970384.

Schaefke, B., Wang, T.-Y., Wang, C.-Y., and Li, W.-H. (2015). Gains and Losses of Transcription Factor Binding Sites in Saccharomyces cerevisiae and Saccharomyces paradoxus. Genome Biol Evol 7, 2245–2257. doi:10.1093/gbe/evv138.

Scharf, D. H., Chankhamjon, P., Scherlach, K., Heinekamp, T., Willing, K., Brakhage, A. A., et al. (2013). Epidithiodiketopiperazine biosynthesis: a four-enzyme cascade converts glutathione conjugates into transannular disulfide bridges. Angew Chem Int Ed Engl 52, 11092–11095. doi:10.1002/anie.201305059.

Scharf, D. H., Remme, N., Habel, A., Chankhamjon, P., Scherlach, K., Heinekamp, T., et al. (2011). A Dedicated Glutathione *S*-Transferase Mediates Carbon–Sulfur Bond Formation in Gliotoxin Biosynthesis. J. Am. Chem. Soc. 133, 12322–12325. doi:10.1021/ja201311d.

Schrettl, M., Carberry, S., Kavanagh, K., Haas, H., Jones, G. W., O’Brien, J., et al. (2010). Self-Protection against Gliotoxin—A Component of the Gliotoxin Biosynthetic Cluster, GliT, Completely Protects Aspergillus fumigatus Against Exogenous Gliotoxin. PLOS Pathogens 6, e1000952. doi:10.1371/journal.ppat.1000952.

Shibuya, K., Takaoka, M., Uchida, K., Wakayama, M., Yamaguchi, H., Takahashi, K., et al. (1999). Histopathology of experimental invasive pulmonary aspergillosis in rats: pathological comparison of pulmonary lesions induced by specific virulent factor deficient mutants. Microb Pathog 27, 123–131. doi:10.1006/mpat.1999.0288.

Shimodaira, H. (2002). An Approximately Unbiased Test of Phylogenetic Tree Selection. Systematic Biology 51, 492–508. doi:10.1080/10635150290069913.

Steenwyk, J. L., Buida, T. J., Labella, A. L., Li, Y., Shen, X.-X., and Rokas, A. (2021a). PhyKIT: a broadly applicable UNIX shell toolkit for processing and analyzing phylogenomic data. Bioinformatics. doi:10.1093/bioinformatics/btab096.

Steenwyk, J. L., Iii, T. J. B., Li, Y., Shen, X.-X., and Rokas, A. (2020a). ClipKIT: A multiple sequence alignment trimming software for accurate phylogenomic inference. PLOS Biology 18, e3001007. doi:10.1371/journal.pbio.3001007.

Steenwyk, J. L., Mead, M. E., de Castro, P. A., Valero, C., Damasio, A., dos Santos, R. A. C., et al. (2021b). Genomic and Phenotypic Analysis of COVID-19-Associated Pulmonary Aspergillosis Isolates of Aspergillus fumigatus. Microbiol Spectr 9. doi:10.1128/Spectrum.00010-21.

Steenwyk, J. L., Mead, M. E., Knowles, S. L., Raja, H. A., Roberts, C. D., Bader, O., et al. (2020b). Variation Among Biosynthetic Gene Clusters, Secondary Metabolite Profiles, and Cards of Virulence Across Aspergillus Species. Genetics 216, 481–497. doi:10.1534/genetics.120.303549.

Steenwyk, J. L., Shen, X.-X., Lind, A. L., Goldman, G. H., and Rokas, A. A Robust Phylogenomic Time Tree for Biotechnologically and Medically Important Fungi in the Genera Aspergillus and Penicillium. mBio 10, e00925–19. doi:10.1128/mBio.00925-19.

Steinbach, W. J., Marr, K. A., Anaissie, E. J., Azie, N., Quan, S.-P., Meier-Kriesche, H.-U., et al. (2012). Clinical epidemiology of 960 patients with invasive aspergillosis from the PATH Alliance registry. Journal of Infection 65, 453–464. doi:10.1016/j.jinf.2012.08.003.

Suyama, M., Torrents, D., and Bork, P. (2006). PAL2NAL: robust conversion of protein sequence alignments into the corresponding codon alignments. Nucleic Acids Research 34, W609–W612. doi:10.1093/nar/gkl315.

Takahashi, H., Umemura, M., Ninomiya, A., Kusuya, Y., Shimizu, M., Urayama, S., et al. (2021). Interspecies Genomic Variation and Transcriptional Activeness of Secondary Metabolism-Related Genes in Aspergillus Section Fumigati. Frontiers in Fungal Biology 2, 14. doi:10.3389/ffunb.2021.656751.

Traynor, A. M., Sheridan, K. J., Jones, G. W., Calera, J. A., and Doyle, S. (2019). Involvement of Sulfur in the Biosynthesis of Essential Metabolites in Pathogenic Fungi of Animals, Particularly Aspergillus spp.: Molecular and Therapeutic Implications. Frontiers in Microbiology 10, 2859. doi:10.3389/fmicb.2019.02859.

Vargas-Muñiz, J. M., Renshaw, H., Richards, A. D., Lamoth, F., Soderblom, E. J., Moseley, M. A., et al. (2015). The Aspergillus fumigatus septins play pleiotropic roles in septation, conidiation, and cell wall stress, but are dispensable for virulence. Fungal Genet Biol 81, 41–51. doi:10.1016/j.fgb.2015.05.014.

Vinet, L., and Zhedanov, A. (2011). A ‘missing’ family of classical orthogonal polynomials. J. Phys. A: Math. Theor. 44, 085201. doi:10.1088/1751-8113/44/8/085201.

Waddell, P. J., and Steel, M. A. (1997). General time-reversible distances with unequal rates across sites: mixing gamma and inverse Gaussian distributions with invariant sites. Mol Phylogenet Evol 8, 398–414. doi:10.1006/mpev.1997.0452.

Wiesner, D. L., and Klein, B. S. (2017). Lung epithelium: barrier immunity to inhaled fungi and driver of fungal-associated allergic asthma. Curr Opin Microbiol 40, 8–13. doi:10.1016/j.mib.2017.10.007.

Willger, S. D., Grahl, N., and Cramer, R. A. (2009). Aspergillus fumigatus metabolism: clues to mechanisms of in vivo fungal growth and virulence. Med Mycol 47 Suppl 1, S72–79. doi:10.1080/13693780802455313.

Willger, S. D., Puttikamonkul, S., Kim, K.-H., Burritt, J. B., Grahl, N., Metzler, L. J., et al. (2008). A Sterol-Regulatory Element Binding Protein Is Required for Cell Polarity, Hypoxia Adaptation, Azole Drug Resistance, and Virulence in Aspergillus fumigatus. PLOS Pathogens 4, e1000200. doi:10.1371/journal.ppat.1000200.

Wirmann, L., Ross, B., Reimann, O., Steinmann, J., and Rath, P.-M. (2018). Airborne Aspergillus fumigatus spore concentration during demolition of a building on a hospital site, and patient risk determination for invasive aspergillosis including azole resistance. J Hosp Infect 100, e91–e97. doi:10.1016/j.jhin.2018.07.030.

Yamamoto, N., Maeda, Y., Ikeda, A., and Sakurai, H. (2008). Regulation of thermotolerance by stress-induced transcription factors in Saccharomyces cerevisiae. Eukaryot Cell 7, 783–790. doi:10.1128/EC.00029-08.

Yang, Z. (2007). PAML 4: Phylogenetic Analysis by Maximum Likelihood. Molecular Biology and Evolution 24, 1586–1591. doi:10.1093/molbev/msm088.

Zhao, W., Panepinto, J. C., Fortwendel, J. R., Fox, L., Oliver, B. G., Askew, D. S., et al. (2006). Deletion of the regulatory subunit of protein kinase A in Aspergillus fumigatus alters morphology, sensitivity to oxidative damage, and virulence. Infect Immun 74, 4865–4874. doi:10.1128/IAI.00565-06.

