## Supplementary Figures for "Extensive sequence divergence of non-coding regions between *Aspergillus fumigatus*, a major fungal pathogen of humans, and its relatives"

**Supplemental Figures and Figure Legends**

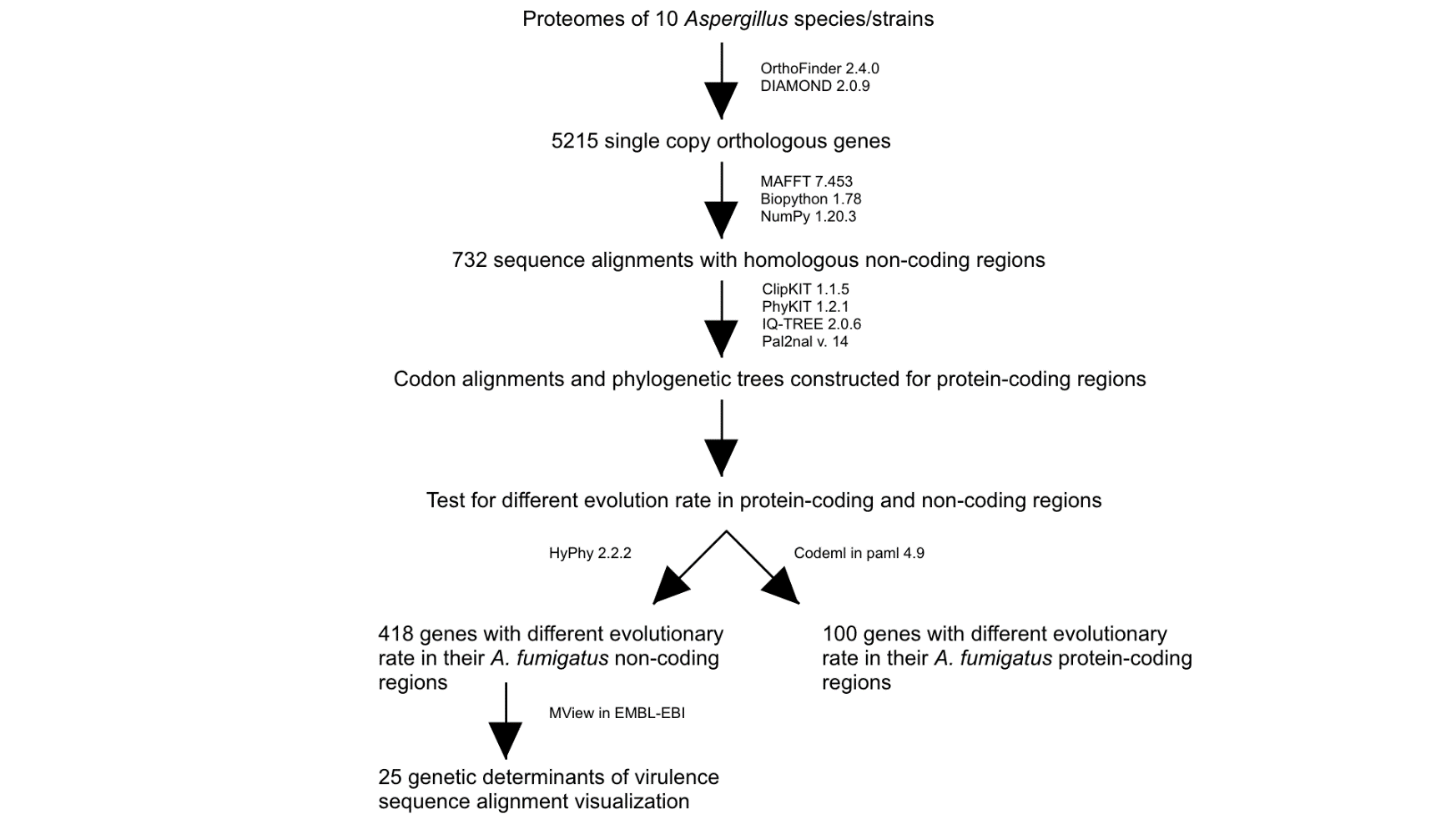

**Supplementary Figure 1. Methods Workflow.** A general outline of the steps taken to go from proteomes of 10 *Aspergillus* species to individual non-coding sequences of interest along with programs used to achieve each step of the pipeline.

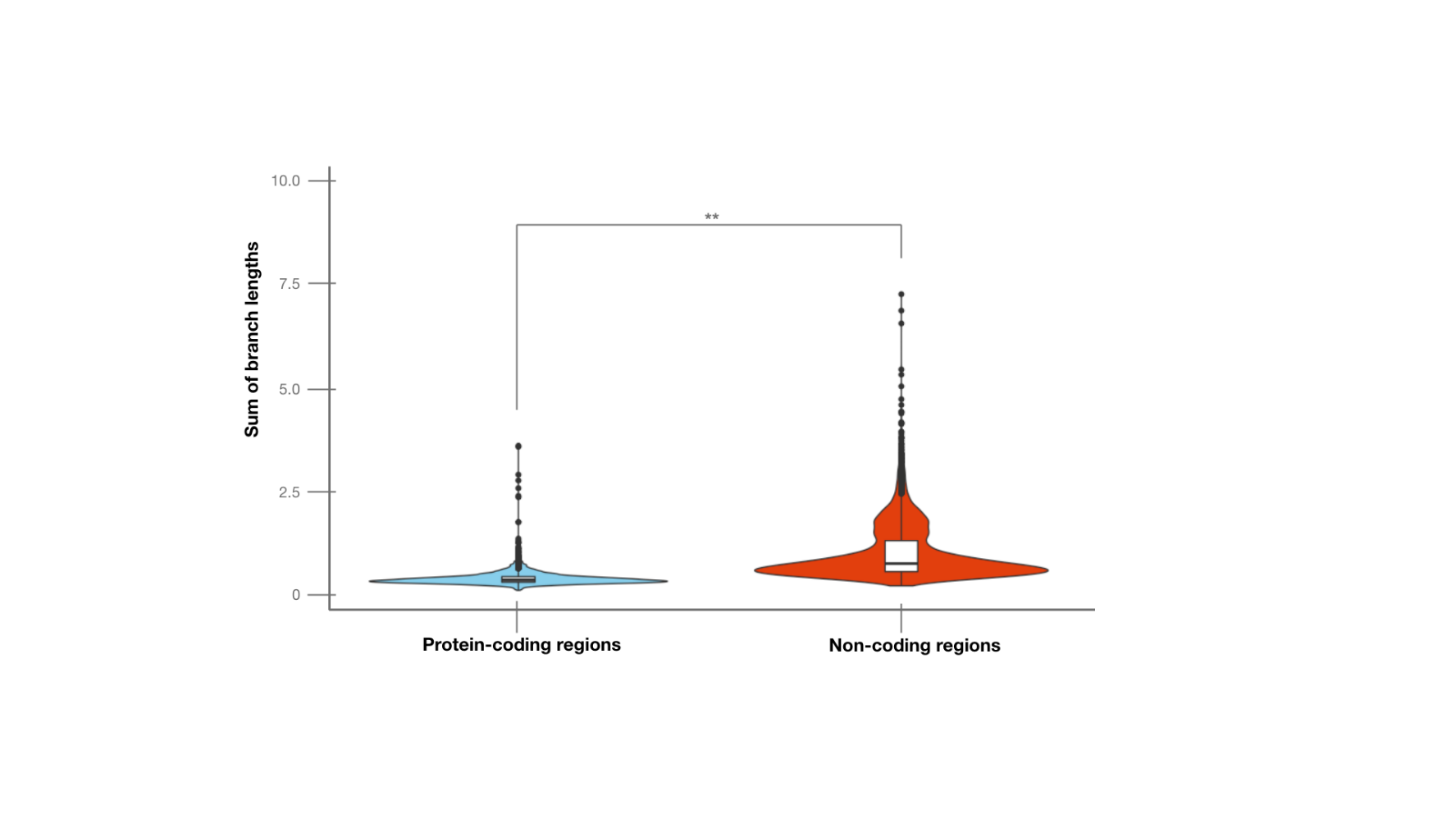

**Supplementary Figure 2. Tree Branch Lengths.** Distribution of the sum of branch lengths in gene trees from protein-coding and non-coding regions. In blue, violin plot of the sum of branch lengths for the protein-coding region trees of 5,215 single copy orthologous genes. In red, violin plot of the sum of branch lengths for non-coding region trees across 5,215 single copy orthologous genes. Non-coding region trees exhibit a significantly larger average sum of branch lengths compared to protein-coding region trees (Wilcoxon signed-ranked test; p-value = 0.004), suggesting that non-coding regions of single-copy orthologs evolve faster than protein-coding regions.
